# Deep immune profiling pinpoints cellular and molecular drivers of lupus immunopathology

**DOI:** 10.64898/2025.12.08.693110

**Authors:** Masahiro Nakano, Michihiro Kono, Kenichiro Asahara, Takayuki Katsuyama, Satoshi Kubo, Eri Katsuyama, Yuya Fujita, Takahiro Nishino, Hajime Inokuchi, Takahiro Arakawa, Tsugumi Kawashima, Shohei Noma, Akiko Minowa, Reza Bagherzadeh, Yukiro Matsumoto, Jun Inamo, Haruka Takahashi, Bunki Natsumoto, Xuejun Zhang, Sang-Cheol Bae, Akari Suzuki, Hiroaki Hatano, Chikashi Terao, Yoshiya Tanaka, Yoshinori Matsumoto, Kazuhiko Yamamoto, Kazuyoshi Ishigaki

**Author notes:** Correspondence: Masahiro Nakano, M.D., Ph.D., Kazuyoshi Ishigaki, M.D., Ph.D. These authors contributed equally to this work (Masahiro Nakano, Michihiro Kono).

## Abstract

Systemic lupus erythematosus (SLE) is a complex autoimmune disease with an unknown etiology. To pinpoint new disease-relevant cell states and their molecular profiles, we performed an in-depth investigation of multimodal single-cell datasets comprising ∼2.1 million peripheral blood mononuclear cells from 346 donors. By resolving 123 fine-grained cell states across 27 cell types, we identified previously uncharacterized populations distinctively associated with clinical severity and treatment status, including *GZMK*^+^*GZMH*^+^*HLA-DR*^+^ effector memory CD8^+^ T cells (double-positive [DP] EMCD8) and *FOXO1*^+^*ARHGAP15*^+^ T cells. Through extensive statistical frameworks and multimodal approaches, we delineated their aberrant immune signaling networks, transcriptional regulators, key surface proteins, T cell receptor repertoires, and genetic/epigenetic landscapes, underscoring them as candidate drivers of SLE immunopathology. These findings provide new insights into therapeutic target discovery in SLE.

## Introduction

Systemic lupus erythematosus (SLE) is a prototypic autoimmune disease with a broad spectrum of clinical manifestations, triggered by genetic and environmental factors (*1*). Since the pathophysiology of SLE involves various immune cell types, many bulk and single-cell transcriptome studies on peripheral blood and affected organs have been conducted (*2–6*). While these efforts have begun to reveal key gene signatures and core cell populations of SLE, the fine-resolution pathophysiology remains unresolved, and the treatment options are still limited; to date, only two molecular-targeted drugs have been approved for this disease (*7*, *8*).

To understand the complex biology of SLE and identify therapeutic targets, it is crucial to characterize disease-relevant cell states and their molecular signatures at high resolution. We previously conducted a large-scale bulk transcriptome study across 27 immune cell types, identifying cell-type-specific signatures linked to SLE development and exacerbation (*6*). However, as an inherent limitation of bulk studies, we failed to identify more granular cell states within each cell type. While single-cell RNA sequencing (scRNA-seq) can theoretically resolve hundreds of cell states, previous SLE studies have not fully leveraged this potential, profiling only ∼20 major cell types (*5*) or ∼40 cell clusters (*3*, *4*), thereby lacking deeper mechanistic insights. Given that low-frequency cell populations often play pivotal roles in various diseases (*9–13*)—such as peripheral helper T cells (Tph) and autoimmune-associated B cells (ABCs) in autoimmunity—a comprehensive identification of disease-relevant cell states and their orchestrated networks is essential to elucidating the complex pathophysiology and clinical heterogeneity of SLE. Moreover, most single-cell studies have not formally distinguished between quantitative cellular abundance shifts and qualitative transcriptional changes within specific cell types. Although recent tools such as milo (*14*) and CNA (*15*) enable the detection of continuous quantitative shifts, they are not designed to directly evaluate qualitative changes. Thus, it is important to extend these concepts and construct a statistical framework capable of jointly dissecting these two scenarios. Finally, as previous SLE studies have focused primarily on gene expression (*3–5*), integrating multi-layered information is paramount to uncovering novel therapeutic targets within the complex molecular architecture of the disease.

To address these gaps, we extensively investigated multimodal single-cell data of SLE with a total of ∼2.1 million cells from 346 donors (<u>Lu</u>pus-derived <u>P</u>eripheral immune cell <u>In</u>tegrated panels [LuPIN]) to pinpoint granular disease-relevant cell states (**Fig. 1**). Using fine-grained clustering and rigorous validation, we defined 123 cell states across 27 immune cell types. Notably, we identified several previously uncharacterized cell states specifically tracking with severe clinical phenotypes and treatment response. Our statistical framework, dissecting both quantitative and qualitative gene signatures, indicated that key immunological pathways and SLE risk variants were closely linked to cell-state compositional shifts within each cell type. Furthermore, these newly identified cell populations were predicted to engage in intricate immune crosstalk with established pathogenic cell states, potentially driving SLE pathogenesis. Finally, in-depth characterization delineated the multifaceted molecular landscapes of these populations, including transcriptional regulators, epigenetic features, surface proteins, and T-cell receptor (TCR) repertoires. Overall, our intensive approach elucidates the complex immunopathology of SLE, providing a foundation for novel therapeutic strategies.

**Fig. 1.**
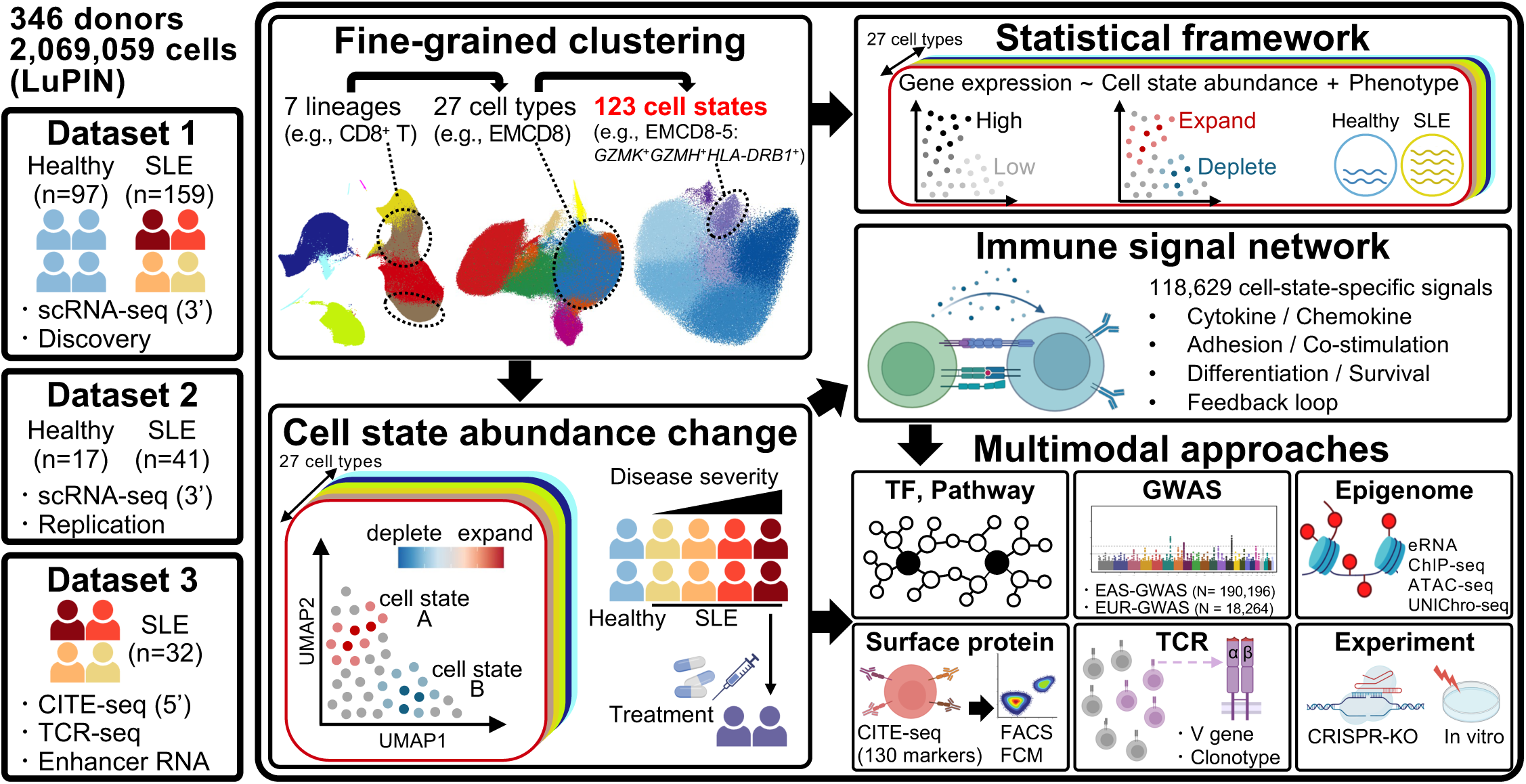
Overview of this study. **(left)** We extensively investigated multimodal single-cell data of SLE of ∼2.1 million cells from 346 donors to pinpoint granular disease-relevant cell states. **(middle)** Using fine-grained clustering, we defined 123 cell states across 27 immune cell types, identifying previously uncharacterized cell states associated with clinical severity and treatment status. **(right)** Our statistical framework indicated that key immunological pathways and SLE risk variants were linked to cell-state compositional shifts. Furthermore, these newly identified cell populations were predicted to engage in immune crosstalk with established pathogenic cell states, alongside in-depth multimodal characterization of their molecular profiles. CITE-seq, cellular indexing of transcriptomes and epitopes by sequencing; TCR, T cell receptor; EMCD8, effector memory CD8^+^ T cells; TF, transcription factor; GWAS, genome-wide association studies; eRNA, enhancer RNA; UNIChro-seq, unique molecular identifier counting of regional chromatin accessibility with sequencing; FCM, flowcytometry; FACS, fluorescence-activated cell sorting.

## Results

### Overview of this study

LuPIN comprises three datasets, each with a different purpose (**Fig. 1, left**). To robustly identify disease-relevant cell states, we first used the largest, publicly available 3’-end scRNA-seq dataset for SLE (*5*) (dataset 1; **table S1**). Our stringent multi-step quality control (QC) process yielded the final dataset (1,456,041 peripheral blood mononuclear cells [PBMCs] from 97 healthy and 172 SLE samples [159 unique SLE donors: 11 donors have 2∼3 timepoints]; **fig. S1**). Among 172 SLE samples, 21 exhibited high disease activity (HDA), and 40 were clinically inactive (**Methods**). Following our previous study scheme (*6*), we established two comparison axes: (i) disease-state comparison between healthy controls and inactive SLE patients, reflecting the biology of disease development, and (ii) disease-activity comparison between inactive vs. HDA SLE, reflecting the biology of disease exacerbation. For replication analysis, we used another scRNA-seq dataset (*4*) (dataset 2, 344,235 PBMCs from 17 healthy and 41 SLE samples).

To overlay in-depth molecular information onto the fine-grained cell state map, we additionally prepared a new multimodal 5’-end dataset using cellular indexing of transcriptomes and epitopes by sequencing (CITE-seq)—a technology profiling gene expression and surface marker proteins from single cells (*16*)—alongside TCR-seq and enhancer RNA profiling (*17*, *18*) (dataset 3, 268,783 PBMCs from 32 SLE samples). Through the efficient integration with dataset 1, we searched for key surface proteins, TCR repertoires, and cis-regulatory elements specific to disease-relevant cell states.

### Definition of 123 robust cell states using fine-grained clustering

To identify granular cell states of SLE, we employed a three-step intensive clustering approach in dataset 1: lineage (layer 1), cell type (layer 2), and cell state (layer 3) (**Fig. 2**). First, Louvain clustering categorized 1,456,041 PBMCs into seven cell lineages (layer 1: CD4^+^ T cells [CD4 T], CD8^+^ and unconventional T cells [CD8 and other T], natural killer cells [NK], B cells, monocytes [Mono], dendritic cells [DC], and hematopoietic stem and progenitor cells [HSPC]; **fig. S1D–F**). These lineages were further subdivided into 27 coarse cell types (layer 2: **Fig. 2A–F**, **fig. S2A–F**, and **table S2**). Our analytical pipelines (**table S3** and **Methods**) achieved higher annotation accuracy than the original study(*5*) (**fig. S2G**).

**Fig. 2.**
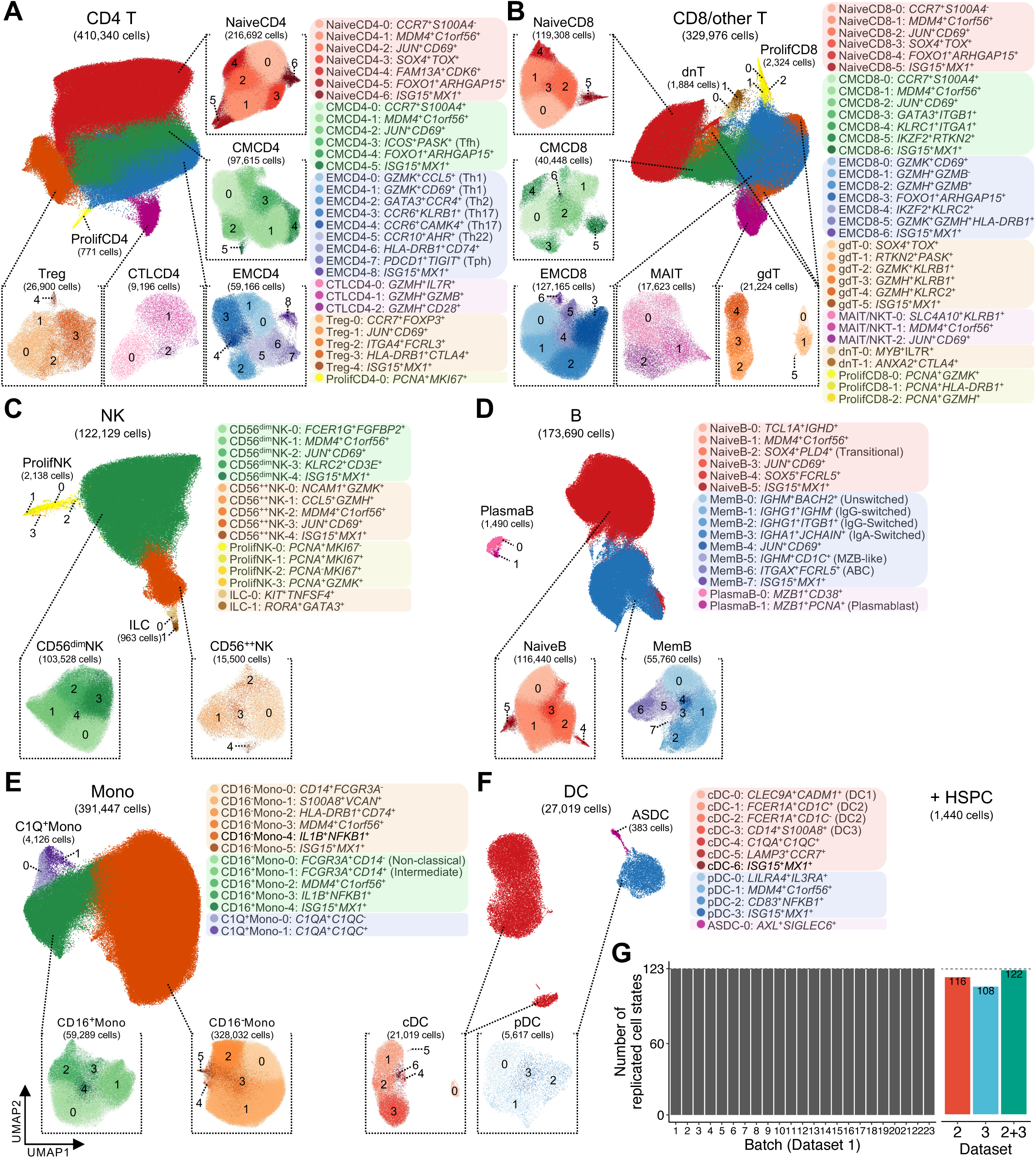
Definition of 123 robust cell states using fine-grained clustering. **(A-F)** UMAP plots showing 27 cell types and 123 cell states identified by fine-grained clustering in **(A)** CD4^+^ T cell (CD4 T), **(B)** CD8^+^ and unconventional T cell (CD8/other T), **(C)** natural killer cell (NK), **(D)** B cell, **(E)** monocyte (Mono), and **(F)** dendritic cell (DC) lineages (dataset 1; *N* = 269). The hematopoietic stem and progenitor cell (HSPC) lineage is omitted as it comprises a single cell type and cell state. For each panel, large lineage-level UMAPs are colored by cell types, while small cell-type-level UMAPs are colored and numbered by cell states. Cell-state annotations are listed to the right of each panel. See **table S2** for cell type abbreviations. **(G)** Bar plots showing the number of cell states replicated in **(left)** leave-one-batch-out analysis of dataset 1 and **(right)** independent clustering of dataset 2 and 3.

To determine if these coarse cell-type-level annotations sufficiently explain disease-relevant compositional changes, we performed co-varying neighborhood analysis (CNA) to identify single-cell-resolution transcriptional neighborhoods associated with clinical phenotypes (*15*). We applied CNA to six cell lineages (excluding HSPC) and examined the abundance of 26 coarse cell types (lineage-level CNA; **fig. S2H**). The extent of expansion or depletion for each cell type was quantified at the single-cell level as “neighborhood correlations”. Using two comparison axes—disease-state (inactive SLE vs. healthy, reflecting disease development) and disease-activity CNA (HDA vs. inactive SLE, reflecting disease exacerbation) (*6*)—we recapitulated the depletion of NaiveCD4 and mucosal-associated invariant T cells (MAIT) in disease-state comparison (*5*), while revealing that these patterns were remarkable in disease-activity comparison. Similarly, regulatory T cells (Treg), proliferating (Prolif) CD4/CD8/NK cells, and C1Q^+^ monocytes (Mono) expanded in disease-state and/or -activity CNA. Although neighborhoods in these cell types exhibited universal expansion or depletion patterns, many other cell types showed heterogeneous patterns in lineage-level CNA (**fig. S2H**), underscoring the need for a more fine-grained investigation.

To efficiently capture subtle gene-program heterogeneities within each cell type, we further performed clustering on each cell type and defined 123 cell states with unique gene expression profiles (layer 3: **Fig. 2A–F**, **fig. S3**, and **table S4**). As these cell-state definitions form the basis for downstream analyses, we employed multiple rigorous approaches to ensure their robustness. First, a leave-one-batch-out analysis, systematically excluding each of the 23 experimental batches from dataset 1, confirmed the consistent preservation of all 123 cell states, albeit occasionally requiring slightly higher resolution parameters (**Fig. 2G, left**, **fig. S4A**, and **table S5**). Moreover, 122 cell states (except for gamma-delta T cells [gdT]-5: *ISG15*^+^*MX1*^+^) were replicated through independent clustering of datasets 2 and/or 3, demonstrating the cross-dataset reproducibility (**Fig. 2G, right**, and **fig. S4B–D**). Second, the identity of each cluster was corroborated by an extensive literature review (*9–13*, *19–39*) (**table S6**). Third, marker genes upregulated in each state exhibited minimal overlap within the same cell type (**fig. S4E**), and pathway analysis using these markers revealed enrichment of cell-state-specific pathways (**table S7**), supporting their non-redundancy and biological relevance. Finally, multimodal investigations using dataset 3 identified cell-state-specific surface proteins and TCR features (**Fig. 6–7**), providing orthogonal validation at the immune phenotypic level.

Our fine-grained annotations successfully captured multiple disease-relevant cell state candidates (**Fig. 2**), most of which remained unidentified in the original study due to insufficient resolution. We first recapitulated established pathogenic states including: (i) follicular and peripheral helper T cells (Tfh and Tph) within central and effector memory CD4 T cells (CMCD4-3: *ICOS*^+^*PASK*^+^ and EMCD4-7: *PDCD1*^+^*TIGIT*^+^) and (ii) autoimmune-associated B cells (ABCs) in memory B cells (MemB-6: *ITGAX*^+^*FCRL5*^+^), which are central to autoantibody production (*9*, *10*, *27*). We also confirmed the presence of subpopulations highly activated by interferon (IFN) signaling (*ISG15*^+^*MX1*^+^ cell states)—a hallmark signature of SLE (*1*, *26*)—across most cell types. The shared gene program across cell types was not limited to IFN signaling; *JUN*^+^*CD69*^+^ cell states with upregulated AP-1 transcription factors (TFs; e.g., *JUN, FOS*) and early activation markers (e.g., *CD69* and *NFKBIA*) (*21*) were identified in most lymphoid cell types (**fig. S3A–D**). Importantly, we identified multiple previously uncharacterized cell states with distinct gene programs, some of which were later found to expand especially in severe SLE (e.g., EMCD8-5: *GZMK*^+^*GZMH*^+^*HLA-DRB1*^+^, **Fig. 3**). Moreover, our approach captured very rare subpopulations such as *ANXA2*^+^*CTLA4*^+^ double-negative T cells (dnT-1) (*32*), *RORA*^+^*GATA3*^+^ innate lymphoid cells (ILC-1) (*33*), *MZB1*^+^*PCNA*^+^ plasmablasts (PlasmaB-1) (*35*), and *LAMP3*^+^*CCR7*^+^ DCs (cDC-5) (*39*). Overall, our intensive clustering approach provided high-resolution cell state information for SLE, serving as the foundation for subsequent analyses.

**Fig. 3.**
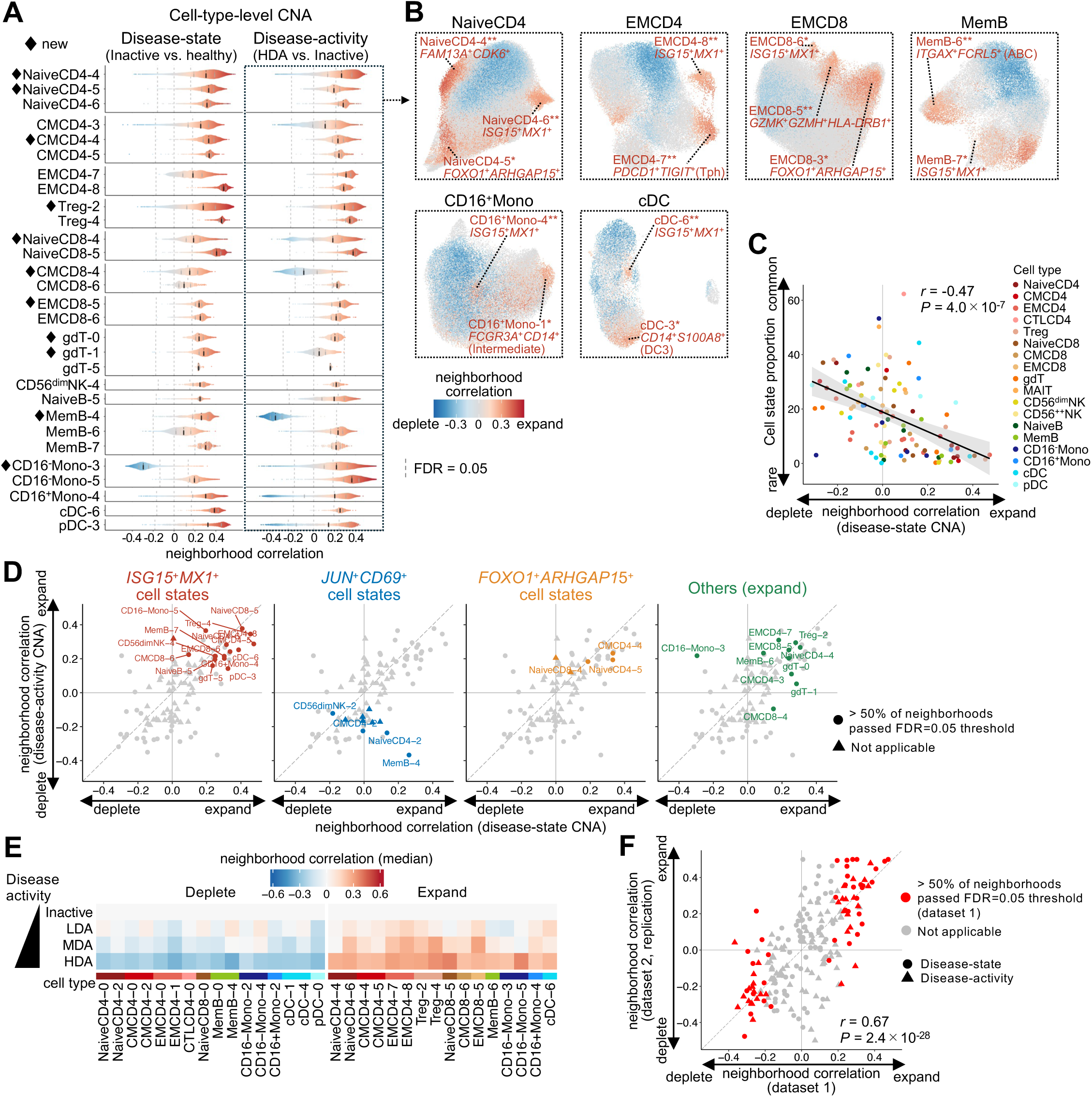
Mapping cell state compositional changes in SLE. **(A)** Distributions of neighborhood correlations for each cell state in disease-state (*n* = 137) or disease-activity (*n* = 61) comparisons (dataset 1, cell-type-level-CNA). 29 cell states with > 50% of neighborhoods positively associated with phenotypes (FDR < 0.05) are shown (See **fig. S5A** for details). **(B)** UMAP plots of representative cell types, colored by neighborhood correlations. Cell states are highlighted if > 50% (**) or > 25% (*) of their neighborhoods are associated with phenotypes. **(C)** Scatter plot of median neighborhood correlation (disease-state CNA) for each cell state vs. cell-state proportion within cell type. Shaded regions represent 95% confidence intervals. **(D)** Comparison of median neighborhood correlation between disease-state (x-axis) and disease-activity (y-axis) CNA. Cell states relevant to each panel title are highlighted in color. **(E)** Heatmap of median neighborhood correlation for low (LDA) / moderate (MDA) / high disease activity (HDA) vs. Inactive SLE (*n* = 58/17/21/40, respectively). 32 disease-activity-associated cell states are shown. **(F)** Comparison of median neighborhood correlation for each cell state between dataset 1 (x-axis) and dataset 2 (y-axis; *N* = 58). In **(C)** and **(F)**, Pearson’s *r* and *P* values are indicated.

### Mapping cell state compositional changes in SLE

To assess the cell-state compositional changes within each cell type, we next applied CNA to 18 major cell types (>5,000 cells; cell-type-level CNA) in disease-state (inactive SLE [*n* = 40] vs. healthy [*n* = 97]) and disease-activity (HDA [*n* = 21] vs. inactive SLE [*n* = 40]) comparisons, accounting for age, sex, and ancestry. Strikingly, we observed skewed neighborhood abundance in 34 out of 36 tests (18 cell types × 2 comparisons, global *P* < 0.05/36), and 29 cell states showed evidence of remarkable expansion (**Fig. 3A–B**, **fig. S5A**, and **table S8**). These expanded cell states tended to be rare within each cell type, supporting the importance of the fine-grained strategy (**Fig. 3C** and **fig. S5B**).

Among the 29 cell states exhibiting remarkable expansion,18 recapitulated known signals. All IFN-activated (*ISG15*^+^*MX1*^+^) cell states within each cell type expanded in disease-state and -activity CNA (**Fig. 3D**). While Tfh (CMCD4-3) expanded primarily in disease-state, Tph (EMCD4-7) expanded specifically in disease-activity CNA. Notably, in memory B cells (MemB)—which harbor the highest enrichment of SLE risk variants (*40*)—ABC (MemB-6) showed remarkable expansion in disease-activity CNA but not in disease-state CNA (**Fig. 3A**), supporting their relevance in pathogenic autoantibody production, particularly during disease activation (*10*). Furthermore, our ancestry-specific analysis revealed the expansion of this population in Asian SLE compared to European (EUR) SLE (**fig. S5C**), suggesting a potential need for stratified medicine using B-cell targeted therapies (*41*, *42*).

The remaining 11 cell states represented previously unreported expansions in SLE (*3–5*), nominating them as new disease-relevant candidates. An example is *GZMK*^+^*GZMH*^+^*HLA-DRB1*^+^ cells (EMCD8-5), which showed expansion in both disease-state and -activity CNA (**Fig. 3A–B**). While other EMCD8 cell states express either *GZMK* or *GZMH*, this population expresses both, along with *GZMB* to a lesser extent (**fig. S3B**; hereafter “double-positive [DP] EMCD8”). Notably, this population exhibited high expression of multiple T cell activation markers, including *HLA-DR* genes, *PDCD1*, and *TIGIT* (**fig. S3B**). Another example, *FAM13A*^+^*CDK6*^+^ cells (NaiveCD4-4), further highlighted the necessity of fine-grained analysis: although NaiveCD4 cells were, on average, depleted within the CD4 T lineage (**fig. S2H**), this depletion pattern was not uniform across NaiveCD4 cell states. Lastly, *FOXO1*^+^*ARHGAP15*^+^ cell states expanded across most T-cell cell types in disease-state and -activity CNA (**Fig. 3A–B, D**). These states were broadly shared across the majority of SLE samples, rather than being derived from a limited number of donors (**fig. S5D**). Some cell states showed different patterns of abundance changes between disease-state and activity CNA. While *JUN*^+^*CD69*^+^ cell states expanded in disease-state CNA, they tended to deplete in disease-activity CNA across most cell types (**Fig. 3D**), suggesting distinct roles of these states in SLE depending on the disease phase (*6*).

Our CNA results demonstrated high reproducibility in multiple approaches. First, cluster-based abundance testing (MASC) (*11*) yielded concordant results with CNA (**fig. S5E**). Second, most CNA signals were globally consistent between Asian and EUR ancestries (**fig. S5F**). Third, by independently applying CNA to low (LDA) and moderate disease activity (MDA) cases—both of which were excluded from the discovery set—we observed concordant signals with a gradual increase in disease-activity effects, reflecting the continuous spectrum of SLE activity (**Fig. 3E**). Lastly, disease-state and activity CNA using independent dataset 2 well replicated the signals detected in dataset 1 (**Fig. 3F**). Collectively, these results robustly captured cell-state compositional changes linked to disease development and exacerbation of SLE.

### Statistical framework to dissect quantitative and qualitative changes within cell types

Having identified SLE-associated cellular compositional shifts within each cell type, we next introduced a statistical framework to address the limitations of conventional cell-type-level analyses. Cell-type-level analyses define genes driven by either of the following two distinct scenarios as disease-associated differentially expressed genes (DEGs, **Fig. 4A**): (i) a gene is specifically expressed in a particular cell state whose abundance shifts with phenotype (quantitative change) or (ii) a gene’s per-cell expression is directly dysregulated by the phenotype, regardless of cell-state abundance (qualitative change). To dissect their respective pathogenic effects, our generalized linear mixed-effect model jointly regresses each single-cell expression by (i) a phenotype-driven cell-state abundance term and (ii) a direct phenotype term (e.g., inactive SLE vs. healthy) capturing per-cell gene dysregulation, while adjusting for donor/batch effects and other covariates (**Fig. 4B**; **Methods**). Specifically, we incorporated single-cell-level neighborhood correlations from cell-type-level CNA (**Fig. 3**) into the model as metrics for cell-state abundance changes. By applying this model across 18 major cell types, we identified (i) “abundance signature genes” (quantitative changes) and (ii) “dysregulated signature genes” (qualitative changes, **table S9**). Notably, quantitative changes drove approximately 50% of the variation in the top 1,000 pseudobulk DEGs for each cell type (**fig. S6A**).

**Fig. 4.**
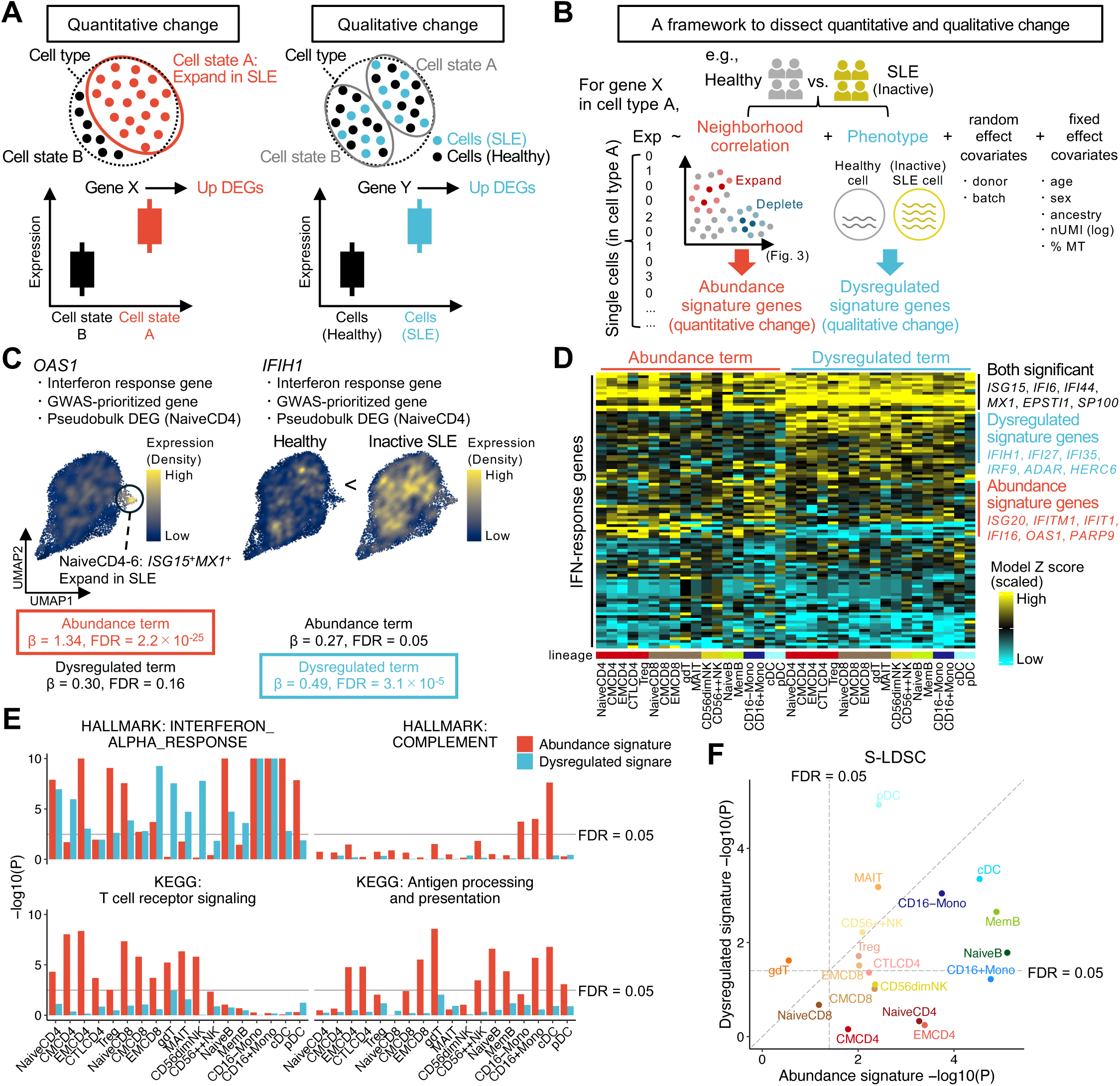
Statistical framework to dissect quantitative and qualitative changes within cell types. **(A)** Schematic representation of two distinct scenarios (quantitative and qualitative changes) driving cell-type-level pseudobulk DEGs. **(B)** Schematic of the framework to dissect quantitative and qualitative changes. **(C)** Representative examples of abundance (*OAS1*) and dysregulated (*IFIH1*) signature genes. UMAPs are colored by expression density. Model statistics for both genes are provided below. **(D)** Heatmap of abundance and dysregulated term Z scores across 18 major cell types for 100 IFN-response genes (dataset 1; disease-state comparison; *n* = 137). Gene order is based on hierarchical clustering of Z scores. **(E)** Bar plots showing the enrichment of representative pathways for each signature across cell types. *P*, *P* values from one-sided Fisher’s exact tests. **(F)** Comparison of SLE risk variant enrichment around abundance vs. dysregulated signature genes for each cell type using stratified LD score regression (S-LDSC).

We tested the robustness of this model through multiple approaches. First, permutation tests showed that this model is well calibrated (**fig. S6B**). Second, a leave-one-batch-out analysis, systematically excluding each batch from dataset 1, confirmed the stability of model statistics across genes (**fig. S6C**), demonstrating the robustness to technical confounders and donor-specific effects. Finally, application of this model to dataset 2 showed good reproducibility of model statistics (**fig. S6D**). For simplicity, we hereafter present disease-state comparison results in **Fig. 4C–F** and disease-activity results in **fig. S6E–F**.

*IFIH1* and *OAS1* are IFN response genes (IRGs) linked to the genetic loci in genome-wide association studies of SLE (*43*) (SLE-GWAS; **table S10**). While cell-type-level pseudobulk analysis identified both as DEGs in NaiveCD4 (false discovery rate [FDR] < 0.05), our model revealed different behaviors of qualitative and quantitative changes for these genes (**Fig. 4C**). While *OAS1* expression localized to NaiveCD4-7 (*ISG15*^+^*MX1*^+^), which expanded in SLE (abundance signature gene), *IFIH1* did not show such localization and was globally overexpressed in SLE compared with healthy cells (dysregulated signature gene). Hierarchical clustering of 100 IRGs (**table S11**) based on model statistics across 18 cell types identified distinct modules representing abundance and dysregulated signature genes, respectively, suggesting heterogeneous roles for IRGs in SLE pathophysiology (**Fig. 4D** and **fig. S6E**).

Next, we performed pathway enrichment analyses on these signatures for each cell type (**Fig. 4E**). IFN signaling was enriched in both dysregulated and abundance signatures across almost all cell types (FDR < 0.05; one-sided Fisher’s exact test), demonstrating that IFN signaling exerts pathogenic effects through both quantitative and qualitative changes. In contrast, other classical immunological pathways were primarily enriched in abundance signature genes of relevant cell types (e.g., complement signaling in myeloid cells; **Fig. 4E**, **fig. S6F**, and **table S12**), suggesting that key biological processes in SLE are more closely linked to abundance signatures than dysregulated signatures.

Finally, we integrated these model statistics with the largest-scale SLE-GWASs in East Asian (EAS) (*44*) and EUR (*45*). We evaluated the polygenic enrichment of SLE risk variants around signature genes using stratified linkage disequilibrium score regression (S-LDSC) (*40*), followed by a meta-analysis of the EAS and EUR results **(fig. S6G**). While SLE risk variants were enriched around both dysregulated and abundance signature genes across multiple cell types (FDR < 0.05), the enrichment was more pronounced for abundance signatures (*P* = 5.3 × 10^−4^; paired Wilcoxon test; **Fig. 4F**). Notably, this enrichment was specifically observed in abundance signature genes with positive statistics (**fig. S6H**). Consistent with previous studies (*40*), strongest enrichment was identified in B cells. Although these findings do not imply that SLE risk variants directly regulate cell-state compositional changes, our framework suggests that SLE risk genes exert their effects primarily within these expanded cell states.

### Characterization of aberrant immune networks between disease-relevant cell states

Motivated by our finding that key immunological pathways and SLE risk variants were predominantly enriched in cell-state compositional changes (**Fig. 4**), we further investigated their clinical relevance. Hierarchical clustering of all samples and cell states based on their proportions revealed that samples broadly partitioned into SLE (sample group [SG]1) and healthy controls (SG2) (**Fig. 5A**). While a subset of SLE patients clustered within SG2, SG1 was further stratified into four subgroups (SG1-1 to SG1-4). Similarly, clustering of cell states identified six cell-state groups (CGs). Intriguingly, while all IFN-activated (*ISG15*^+^*MX1*^+^) cell states correlated within CG3, other SLE-expanded states—including well-known pathogenic (Tfh, Tph, and ABCs) and novel populations (e.g., DP EMCD8 and *FOXO1*^+^*ARHGAP15*^+^ cells)—formed another distinct group (CG2). This suggests that these populations are immunologically interconnected, partially independent of IFN signaling. Indeed, SG1-1 was characterized by a remarkable expansion of CG3, while SG1-2 (and a subset of SG1-3) exhibited CG2 expansion, highlighting the cellular heterogeneity underlying SLE. Both SG1-1 and SG1-2 comprised a higher proportion of HDA patients; conversely, SG1-4 and SG2 were enriched for inactive or LDA patients (**Fig. 5B**). SG1-4 showed *JUN*^+^*CD69*^+^ (CG6) expansion (**Fig. 5A**), potentially marking a clinically quiescent state. We found no significant associations between SGs and organ activity or treatment agents (**fig. S7A**).

**Fig. 5.**
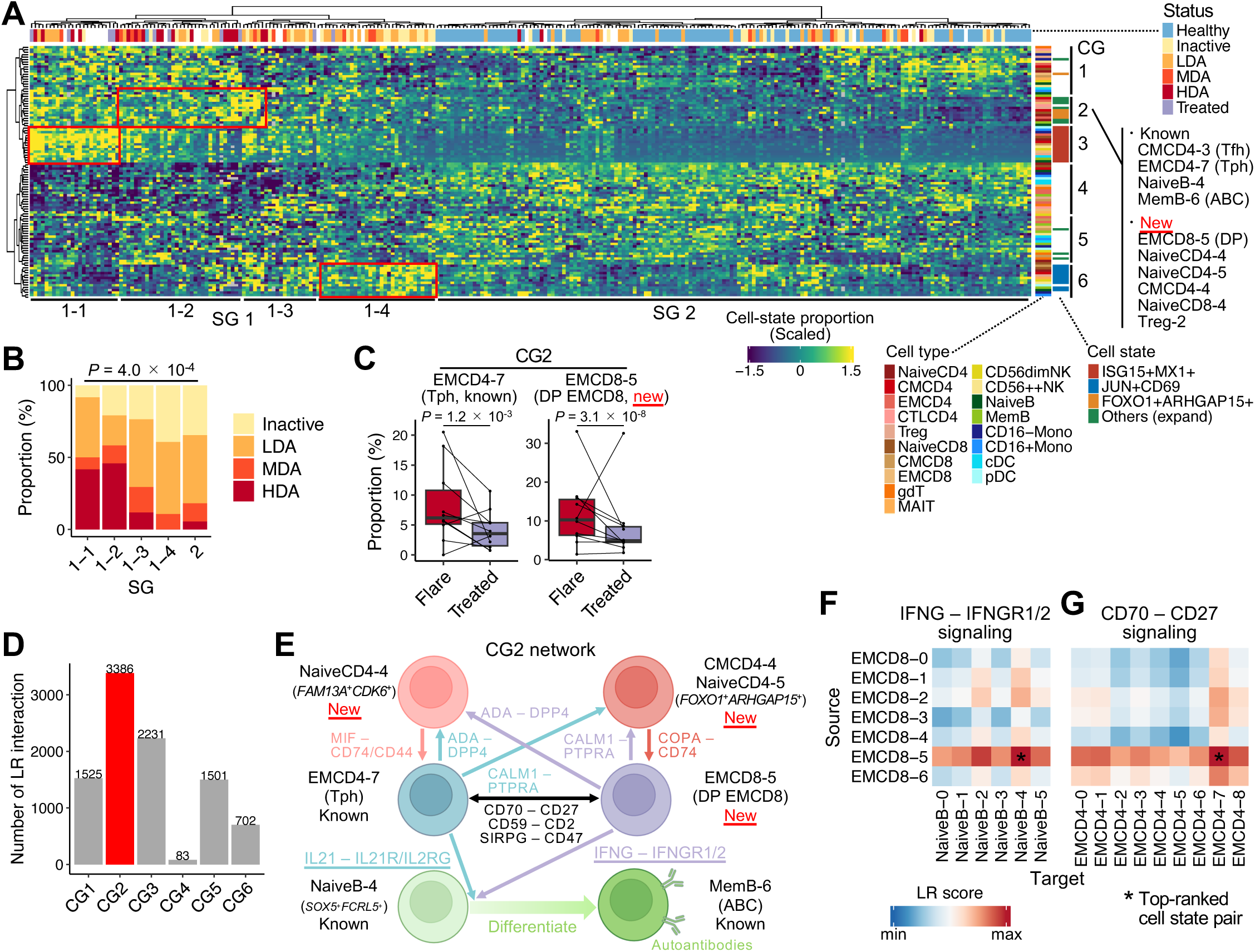
Characterization of aberrant immune networks between disease-relevant cell states. **(A)** Heatmap of cell-state proportions across 18 major cell types for all samples (dataset 1, *N* = 269). Sample and cell-state orders are based on hierarchical clustering of cell-state proportions. **(B)** Bar plot showing the frequencies of disease-activity categories for each sample group (SG). *P*, *P* value from two-sided Fisher’s exact test. LDA/MDA/HDA; low/moderate/high disease activity. **(C)** Box plots comparing cell-state proportions between “Flare” and “Treated” status in **(left)** Tph and **(right)** DP EMCD8. Samples from the same donor are connected by lines. *P*, *P* values from MASC (*n* = 10×2). **(D)** Bar plot showing the number of potential cell-state-specific ligand-receptor (LR) interactions within each CG. **(E)** Schematic of cell-state-specific LR networks across known pathogenic and newly identified population in cell-state group (CG) 2. **(F-G)** Heatmaps of LR scores for **(F)** IFNG–IFNGR1/2 signaling from EMCD8 to NaiveB cell states and **(G)** CD70–CD27 signaling from EMCD8 to EMCD4 cell states.

**Fig. 6.**
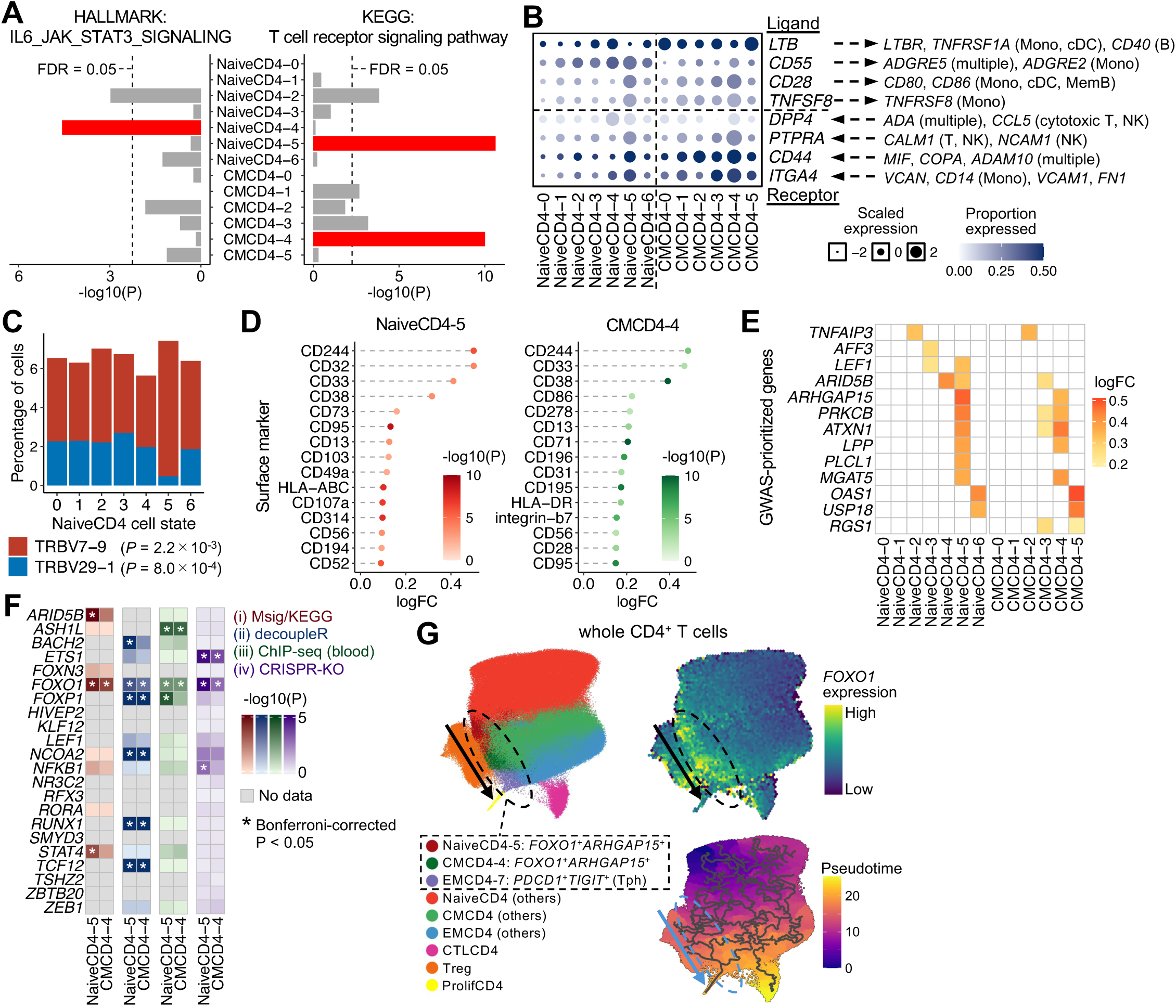
*FOXO1* is a candidate driver of the pathogenic CD4^+^ T cell state gene program. **(A)** Bar plots showing the enrichment of representative pathways for upregulated signature genes for NaiveCD4 and CMCD4 cell states. *P*, *P* values from one-sided Fisher’s exact tests. **(B)** Dot plot showing scaled expression levels of ligand and receptor genes across NaiveCD4 and CMCD4 cell states. Color intensity represents the proportion of cells expressing each gene. Corresponding interaction partners are indicated on the right. **(C)** Bar plot showing the percentage of two TCR V gene usages skewed in NaiveCD4-5. *P*, *P* values from two-sided Fisher’s exact tests. **(D)** Dot plot showing the logFC of top 15 upregulated surface markers for NaiveCD4-5 and CMCD4-4. *P*, *P* values from Seurat FindMarkers. **(E)** Heatmap showing the logFC of GWAS-prioritized genes across NaiveCD4 and CMCD4 cell states. Up to five genes with the highest logFC per cell state are shown. **(F)** Heatmap showing - log_10_(enrichment *P*) values of each TF for NaiveCD4-5 or CMCD4-4 signature genes based on (i) pathway/TF enrichment analyses, TF activity inference using (ii) decoupleR and (iii) blood-derived ChIP-seq data, and (iv) an arrayed CRISPR-KO experiment. *P*, *P* values from one-sided Fisher’s exact tests. **(G)** UMAP plots of whole CD4^+^ T cells, colored by **(left)** focused cell states and other cell types, **(upper right)** *FOXO1* expression, and **(lower right)** pseudotime from trajectory analysis. Edges represent trajectory graph.

**Fig. 7.**
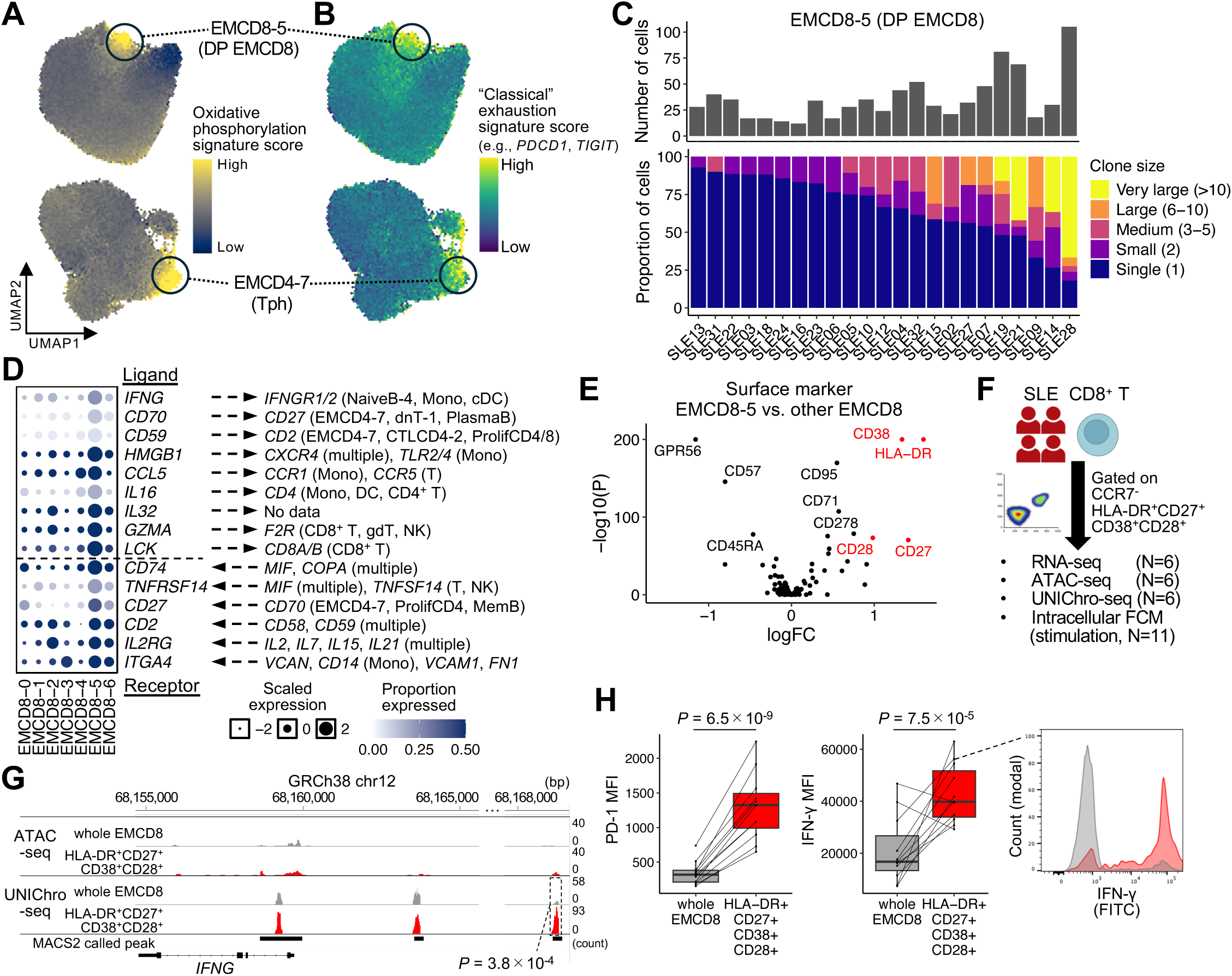
DP EMCD8 cells as a candidate pathogenic player in SLE. **(A-B)** UMAP plots of **(top)** EMCD8 and **(bottom)** EMCD4, colored by **(A)** oxidative phosphorylation and **(B)** exhaustion signature scores. **(C)** Bar plots showing **(top)** the absolute number of EMCD8-5 cells per donor and **(bottom)** the proportion of TCR clonotypes categorized into five size groups within EMCD8-5 (dataset 3; *N* = 32; 23 donors with >10 cells are shown). **(D)** Dot plot showing scaled expression levels of ligand and receptor genes across EMCD8 cell states. Color intensity represents the proportion of cells expressing each gene. Corresponding interaction partners are indicated on the right. **(E)** Volcano plot showing differential expression of surface proteins between EMCD8-5 vs. other EMCD8 cells. *P*, *P* values from Seurat FindMarkers. **(F)** Experimental design for flowcytometry (FCM) and FACS of HLA-DR^+^CD27^+^CD38^+^CD28^+^ EMCD8 cells. **(G)** ATAC-seq and UNIChro-seq tracks around *IFNG* locus in whole EMCD8 and HLA-DR^+^CD27^+^CD38^+^CD28^+^ cells from an SLE donor. *P*, logistic regression *P* values. **(H) (left)** Boxplots comparing the mean fluorescence intensity (MFI) of PD-1 and IFN-γ between HLA-DR^+^CD27^+^CD38^+^CD28^+^ and whole EMCD8 cells. Samples from the same donor are connected by lines. *P*, *P* values from linear mixed models. **(right)** Representative histogram comparing IFN-γ fluorescence intensity between HLA-DR^+^CD27^+^CD38^+^CD28^+^ and whole EMCD8 cells from an SLE donor.

Leveraging longitudinal samples in dataset 1 (from “Flare” to “Treated” samples from 10 donors), we compared changes in cell-state proportions. Notably, while many CG2 cell states (e.g., Tph and DP EMCD8) decreased following treatment, CG3 (*ISG15*^+^*MX1*^+^) remained stable or even increased (**Fig. 5C** and **fig. S7B**), suggesting that CG2 might serve as a cellular biomarker reflecting disease activity and treatment response.

Next, we inferred the inflammatory networks underlying these correlated cell states. By evaluating 4,701 ligand**–**receptor (LR) interactions (*46*) across all 123 × 123 cell-state pairs, we screened an extensive signaling space and identified 1,451,770 potentially active signals (LR score > 0.6; **Methods**). To minimize redundancy, we prioritized the most upregulated cell-state pair as the representative for each unique LR interaction within each of the 27 × 27 cell-type pairs. Among these 118,629 cell-state-specific interactions (of which 52,599 occurred within 18 × 18 major cell-type pairs), 9,428 were predicted to occur between states within the same CG. Intriguingly, within-CG2 interactions accounted for more than one-third (3,386) of these signals, suggesting a prominent immune network among SLE-expanded cell states (**Fig. 5D**). Within CG2, we delineated both established and novel candidate disease-relevant networks (**Fig. 5E** and **table S13**). First, our analysis recapitulated enhanced IL21-IL21R/IL2RG signaling from EMCD4-7 (Tph) to NaiveB-4 (*SOX5*^+^*FCRL5*^+^), a signaling axis essential for their differentiation into MemB-6 (ABC) (*10*). Notably, we identified potential IFNG-IFNGR signaling from EMCD8-5 (DP EMCD8) to NaiveB-4, which is also implicated in ABC differentiation (**Fig. 5E–F**), suggesting that Tph and DP EMCD8 may synergistically drive the maturation of pathogenic B cells (*10*, *47*). Furthermore, Tph and DP EMCD8 were predicted to engage in bidirectional costimulatory crosstalk (e.g., CD70-CD27 signaling) (*48*, *49*), mutually enhancing their effector functions (**Fig. 5E, G**). Additionally, other CG2 cell states, including NaiveCD4-4 (*FAM13A*^+^*CDK6*^+^), NaiveCD4-5 and CMCD4-4 (*FOXO1*^+^*ARHGAP15*^+^), were predicted to amplify these pro-inflammatory cascades (*50–52*) (**Fig. 5E** and **fig. S7C–E**). Collectively, high-resolution cell–cell interaction analysis provided a putative map of intricate immune networks between established and newly identified pathogenic cell states, potentially driving the complex immunopathology of SLE.

### Multimodal molecular profiling across fine-grained cell states

To characterize multi-layered molecular profiles across fine-grained cell states, we prepared a new dataset (dataset 3; 32 SLE donors) comprising CITE-seq (130 surface marker proteins), TCR-seq, and 5’ transcript-based enhancer RNA (eRNA) profiling. Reference mapping (*53*) successfully integrated 268,783 PBMCs from dataset 3 into the latent transcriptional space in dataset 1, extending our fine-grained analysis across multimodal layers (**fig. S8A**). First, we comprehensively interrogated cell-state-specific surface protein expression, identifying an average of nine upregulated proteins per cell state (FDR < 0.05; **fig. S8B** and **table S14**). This analysis recapitulated well-established markers, such as (i) CD278 (ICOS), CD279 (PD1), and TIGIT in Tph (EMCD4-7) (*9*), (ii) CD11c and CD19 in ABC (MemB-6) (*10*), and (iii) CD38 and CD31 in *SOX4*^+^ naive T cell states (NaiveCD4-3 and NaiveCD8-3) (*22*). Furthermore, we identified several key proteins characterizing new disease-relevant cell states, which facilitated our subsequent downstream investigations (**Fig. 6–7**).

We next examined TCR repertoire profiles. While we and others have shown that V gene usage differs across cell types (*54*, *55*) and represents a potential therapeutic target for autoimmunity (*56*), systematic analysis at the cell-state resolution has been lacking. Principal component analysis (PCA) of V-gene frequencies confirmed that cell-state embeddings globally clustered by cell type (**fig. S8C**). Notably, among 94 V genes (45 α- and 49 β-chains), ∼3.8 V genes per cell state exhibited skewed usage (FDR < 0.05; two-sided Fisher’s exact test; **fig. S8D** and **table S15**), highlighting distinct TCR signatures unique to specific cell states. While most of them were observed in EMCD8 cell states, we identified several cell-state-specific V genes in other cell types (e.g., NaiveCD4-5; **Fig. 6**).

Leveraging 5’ transcript sequencing, we mapped transcription start sites (TSSs) at single-cell resolution. Recent studies have shown that bidirectional transcription around TSSs marks enhancer activity (*17*, *18*). Using this approach, we defined ∼1,564 transcribed enhancers per cell type (**Methods**), which were enriched for established PBMC enhancer marks (*57*) (**fig. S8E**). Among them, we identified ∼68 cell-state-specific enhancers per cell type (**table S16**). Thus, the efficient integration of multimodal data with large-scale scRNA-seq data provided a comprehensive molecular map of fine-grained cell states.

### *FOXO1* is a candidate driver of the pathogenic CD4^+^ T cell state gene program

We next investigated the molecular profiles of disease-relevant cell state candidates that exhibited expansion in both disease-state and disease-activity CNA (**Fig. 3A–B**). Specifically, we interrogated the molecular features of newly identified CD4^+^ T cell states within CG2 (**Fig. 5**): (1) *FAM13A*^+^*CDK6*^+^ cells (NaiveCD4-4), and (2) *FOXO1*^+^*ARHGAP15*^+^ cells (NaiveCD4-5 and CMCD4-4).

Pathway analysis of upregulated marker genes revealed an enrichment of JAK-STAT signaling in *FAM13A*^+^*CDK6*^+^ cells and TCR signaling in *FOXO1*^+^*ARHGAP15*^+^ cells, reflecting their distinct immunological roles (**Fig. 6A** and **table S7**). Cell-state-specific enhancers near the marker genes suggested unique epigenetic regulations of these states (**fig. S9A** and **table S16**). Regarding ligand-receptor signaling, both cell states were predicted to engage in pro-inflammatory networks within CG2, including ADA–DPP4, CALM1–PTPRA, MIF–CD74/CD44, and COPA–CD74 axes (*50–52*), which interact with Tph and DP EMCD8 cells (**Fig. 5D**, **fig. S7D–F, and table S13**). Beyond the CG2 network, these states exhibited unique ligand-receptor expression profiles, potentially forming extensive crosstalk with multiple cell types (**Fig. 6B**). Consistent with the upregulated TCR signaling pathway, NaiveCD4-5 and CMCD4-4 highly expressed *CD28*, *TNFSF8*, *PTPRA*, *CD44*, and *ITGA4*, suggesting an “effector-primed” states triggered by persistent antigen stimulation (*49*, *50*, *58*); indeed, NaiveCD4-5 further exhibited a unique TCR V gene usage pattern (**Fig. 6C** and **table S15**).

Surface protein profiling further characterized these states: *FAM13A*^+^*CDK6*^+^ cells highly expressed CD127 (IL-7Rα) (FDR < 0.05; **fig. S9B** and **table S14**), reflecting their distinct homeostatic requirements (*59*). In *FOXO1*^+^*ARHGAP15*^+^ cells, the upregulation of specific activation markers including CD244, CD32, CD33, and CD38 was consistent with chronic antigen exposure via TCR signaling (*60*, *61*) (**Fig. 6D**). Intriguingly, these upregulations were less pronounced at the mRNA level (**table S4**), highlighting the significance of multi-layered investigation (*13*, *16*, *19*). We also examined the localization of GWAS-prioritized genes: in *FAM13A*^+^*CDK6*^+^ cells, the transcription coactivator *ARID5B* was linked to an SLE-GWAS locus (*43*) (**Fig. 6E** and **table S10**). In *FOXO1*^+^*ARHGAP15*^+^ cells, multiple genes including *ARHGAP15* and *PRKCB* were associated with SLE-GWAS loci.

To identify transcriptional regulators driving the gene program of *FAM13A*^+^*CDK6*^+^ and *FOXO1*^+^*ARHGAP15*^+^ cells, we integrated computational and experimental approaches, focusing on 22 candidate TFs upregulated in these states: (i) pathway and TF enrichment analyses, TF activity inference based on (ii) decoupleR (*62*) and (iii) curated ChIP-seq data (*63*) limited to blood tissue, and (iv) an arrayed CRISPR-KO experiment in primary CD4^+^ T cells (**fig. S9C** and **table S17–19**). Notably, *FOXO1* consistently showed significant enrichment for NaiveCD4-5 and CMCD4-4 signature genes across all approaches (Bonferroni-corrected *P* < 0.05, **Fig. 6F**). Within EMCD4 cells, *FOXO1* exhibited its highest expression in Tph (EMCD4-7; **Fig. 6G** and **fig. S3A**). Trajectory analysis (*64*) on whole CD4^+^ T cells also suggested a potential transcriptional continuity between *FOXO1*^+^*ARHGAP15*^+^ cells and Tph. Recent studies have established that *FOXO1* is essential for maintaining T cell stemness and memory programming (*24*, *25*). While persistent antigen exposure drives the T-cell activation program in *FOXO1*^+^*ARHGAP15*^+^ cells, *FOXO1* may act as a differentiation gatekeeper, preventing terminal exhaustion and maintaining a long-lived reservoir of pathogenic effectors. In contrast, *ARID5B* did not robustly account for the *FAM13A*^+^*CDK6*^+^ gene program across our approaches (**fig. S9D**), although the limitations of the in vitro context must be considered. Together, our multi-layered investigation pinpointed key regulatory candidates in pathogenic CD4^+^ T cell states of SLE.

### DP EMCD8 cells as a candidate pathogenic player in SLE

Among the new cell states within CG2, we next performed in-depth investigation of DP EMCD8 (EMCD8-5; *GZMK*^+^*GZMH*^+^*HLA-DRB1*^+^). Notably, this population shared several transcriptional signatures with Tph beyond cell lineages. Pathway analysis revealed an enrichment of oxidative phosphorylation signaling in both populations, suggesting shared metabolomic reprogramming (**Fig. 7A** and **table S7**). Furthermore, DP EMCD8 exhibited the highest “classical” exhaustion signature (e.g., *PDCD1* and *TIGIT*, **table S20**) within EMCD8, mirroring the pattern of Tph within EMCD4. (**Fig. 7B**). Similarly, *RGS1*—an SLE GWAS-prioritized gene and exhaustion marker—was upregulated in both (**fig. S10A**). While these signatures were traditionally regarded as markers of T cell exhaustion, recent studies in autoimmunity indicate that they reflect a highly activated state induced by chronic antigen exposure, rather than a dysfunctional state (*9*, *65*). Consistently, DP EMCD8 showed evidence of TCR clonal expansion in multiple SLE donors (**Fig. 7C**), with some expanded clonotypes potentially reactive to cytomegalovirus or Epstein-Barr virus epitopes (**table S21**). Although many clones were shared with other EMCD8 states, nine clonotypes and nine V genes were specific to DP EMCD8 (FDR < 0.05; two-sided Fisher’s exact test; **fig. S10B–C, table S15** and **21**).

As discussed above (**Fig. 5E–G**), DP EMCD8 cells were predicted to engage in pro-inflammatory networks with other pathogenic populations. Specifically, DP EMCD8 and Tph may synergistically drive the maturation of pathogenic B cells (ABCs) via *IFNG* and *IL21* signaling, while potentially maintaining a feed-forward activation loop through bidirectional costimulatory crosstalk (e.g., CD70**–**CD27). Longitudinal analysis also showed that both DP EMCD8 and Tph decreased following treatment (**Fig. 5C**), tracking with disease activity. Furthermore, DP EMCD8 exhibited high expression of pro-inflammatory cytokines (*CCL5*, *IL16*, and *IL32*) and functional markers (*CD74*, *CD2*, and *ITGA4*; **Fig. 7D**), underscoring their activated effector function rather than exhaustion. Intriguingly, while DP EMCD8 cells expressed lower *GZMB* levels than classical cytotoxic states (EMCD8-2; **fig. S3B**), they showed marked *GZMA* upregulation (**Fig. 7D**), suggesting that they might also mediate caspase-independent cytotoxicity (*66*).

To identify key surface proteins, we integrated the multimodal dataset 3 into dataset 1, identifying HLA-DR, CD27, CD38, and CD28 as the most upregulated proteins in DP EMCD8 (**Fig. 7E** and **table S14**). Notably, CD45RA was downregulated, indicating that these cells are distinct from terminally differentiated effector memory (TEMRA) cells (*67*). In-silico gating on HLA-DR^+^CD27^+^CD38^+^CD28^+^ achieved >90% purity for DP EMCD8 (**fig. S10D**), enabling validation via flow cytometry (FCM) and FACS. We sorted target EMCD8 cells (CCR7^−^HLA-DR^+^CD27^+^CD38^+^CD28^+^) and performed bulk RNAseq and ATAC-seq (**Fig. 7F**, **fig. S10E**, and **table S22**), confirming a gene expression profile comparable to DP EMCD8 (**fig. S10F**). Motif analysis for differentially accessible regions (DARs) in this population compared with whole EMCD8 (**table S23**) showed minimal TF enrichment (**fig. S10G**). Combined with the few upregulated TFs in this population (**table S4**), this suggests that the primary driver of DP EMCD8 may be extrinsic humoral signals rather than intrinsic transcriptional reprogramming.

Since no DARs were initially detected near the *IFNG* locus in standard ATAC-seq, we employed UNIChro-seq (*68*) to enrich tagmented DNA fragments in this region (**table S24**). A ∼20-fold enrichment compared with standard ATAC-seq identified a unique DAR (**Fig. 7G**). Consistently, this population exhibited higher IFN-γ protein expression than whole EMCD8 upon in-vitro stimulation, alongside elevated PD-1 and TIGIT levels (**Fig. 7H** and **fig. S10H**), suggesting dysregulation of the IFN-γ axis across multimodal layers. Accumulating evidence suggests that IFN-γ (type II IFN), primarily produced by CD8^+^ T cells, plays a pivotal role in SLE, alongside type I IFN (*69–71*). Our strategy successfully pinpointed DP EMCD8 as the cellular origin of this key cytokine with a pro-inflammatory phenotype, underscoring its role as a candidate pathogenic player in SLE.

## Discussion

In this study, we employed an in-depth investigation strategy to pinpoint disease-relevant cell states and molecules from large-scale single-cell datasets of SLE. Fine-grained clustering resolved 123 cell states across 27 cell types, including previously uncharacterized populations distinctively associated with clinical severity and treatment status. Subsequent multimodal characterization delineated cell-state-specific immune signaling networks, transcriptional regulators, epigenetic features, surface proteins, and TCR repertoires.

While recent studies suggest that granular cell states within each cell type can exhibit continuous plasticity (*72*), our results demonstrate that a fine-grained cell-state definition is essential for a deeper understanding of disease pathophysiology, facilitating: (i) the interpretation of cluster-free cell abundance testing (**Fig. 3**), (ii) patient stratification with high clinical relevance based on cell-state abundance (**Fig. 5**), (iii) the delineation of aberrant immune crosstalk (**Fig. 5**), and (iv) the functional characterization of disease-relevant populations and the identification of their potential therapeutic targets (**Fig. 6–7**). Since disease-relevant cell states are often low-frequency populations easily obscured by coarse clustering, our approach is consistent with studies that successfully identified rare pathogenic subsets (*9*, *10*, *13*). Leveraging an extensive literature review (*9–13*, *19–39*) and rigorous validation, we reconstructed 123 cell states with distinct immunological features from the largest publicly available single-cell PBMC dataset (*5*) (∼1.5 million cells). While recent studies on inflamed tissues in autoimmune diseases have identified critical parenchymal subpopulations (*13*, *73*), the resolution of immune cells in those studies remained limited (∼60 immune cell states from 0.2∼0.4 million cells). To benefit the research community, we have released LuPIN, a high-resolution immune cell reference panel integrated with a comprehensive atlas of immune signaling networks and multimodal features. Researchers can map their datasets onto LuPIN for precise cell-state annotation and mechanistic inference, providing opportunities to uncover new pathophysiology across multiple diseases.

We also proposed a statistical framework to dissect quantitative and qualitative changes linked to disease within each cell type. Although methods like miloDE (*74*) infer differential expression unique to transcriptional neighborhoods, they were not designed to jointly dissect abundance shifts and per-cell gene dysregulation. Distinguishing these scenarios is a critical step in identifying therapeutic targets, as each may play distinct roles in pathophysiology. The significant enrichment of SLE risk variants within expanded cell states further suggests that these populations are pivotal for variant-to-function research (*75*). While our model demonstrated reasonable robustness and reproducibility, its performance relies on the latent space defined by current sparse single-cell gene expression. The distinction between abundance and dysregulated signatures may be further refined as single-cell technologies advance.

The identification of cell-state-specific regulators and surface proteins has significant implications for drug discovery. For instance, given that FOXO1 has been implicated in the efficacy of cancer immunotherapy (*24*, *25*), modulating FOXO1 could represent a novel treatment option for autoimmunity by targeting aberrant “effector-primed” CD4^+^ T cell states in SLE. Regarding surface proteins, current therapeutic agents often target ubiquitous markers (e.g., CD19 and CD20 in B cells) (*41*, *42*), which can deplete normal cells alongside autoreactive ones. Given the recent development of bispecific antibodies (*76*), narrowing the target to disease-relevant cell states based on specific markers could provide more precise and efficient treatment options.

This study has some limitations. First, we have yet to identify the optimal combination of surface markers to purify *FOXO1*^+^*ARHGAP15*^+^ T cells; more comprehensive panels may address this in the future. Second, extremely rare cell states within minor cell types could not be included in several downstream analyses due to limited cell numbers (**Fig. 3–4**); improvements in throughput will likely uncover their contribution to pathogenesis. Third, as this study focused on PBMCs, key tissue-resident populations may have been overlooked; thus, our approach should be extended to inflamed tissues across various diseases. Fourth, our study design lacked in-vivo functional validation of the identified cell states. Collectively, our findings contribute to a deeper understanding of SLE pathophysiology and facilitate the discovery of novel therapeutic targets.

## Acknowledgments

The super-computing resource was provided by Human Genome Center, Institute of Medical Sciences, The University of Tokyo. We appreciate the generosity of the donors for a RIKEN solicited donation project, Support for SLE research and development. We appreciate the RIKEN-IMS Genome Platform for its help in the sequencing experiments. We thank AIDA consortium and RIKEN-IMS Laboratory for Gene Structure and Regulation for helpful feedback.

## Authors contributions

M.N. and K.I. designed the study. M.N., K.I., and M.K. wrote the manuscript with critical inputs from T.N., Y.T., and K.Y. M.N. conducted analyses with support from K.I., M.K., K.A., H.I., T.N., J.I., H.T., and H.H. M.K. M.N., and K.A. conducted experiments with support from T. A., T. K., S. N., R. B., Y.M., B.N., A.S., and K.Y. T.K., S.K., E.K., Y.F., Y.T., and Y.M. contributed to sample collection. X.Z., S.B. and C.T. managed EAS SLE-GWAS data. All authors contributed to the final manuscript.

## Data, code, and materials availability

Single-cell count matrices of gene expression, surface protein, and TCR features in the CITE-seq dataset will be available at the National Bioscience Database Center (NBDC) Human Database (https://humandbs.dbcls.jp/en/) as of the date of publication. Public data used in this study are available at https://cellxgene.cziscience.com/collections. Reference panels for each lineage and each cell type are available on GitHub (https://github.com/MasahiroNakano-hub/LuPIN). Original codes used in this study are available on GitHub (https://github.com/MasahiroNakano-hub/LuPIN).

---

**Fig. S1.**
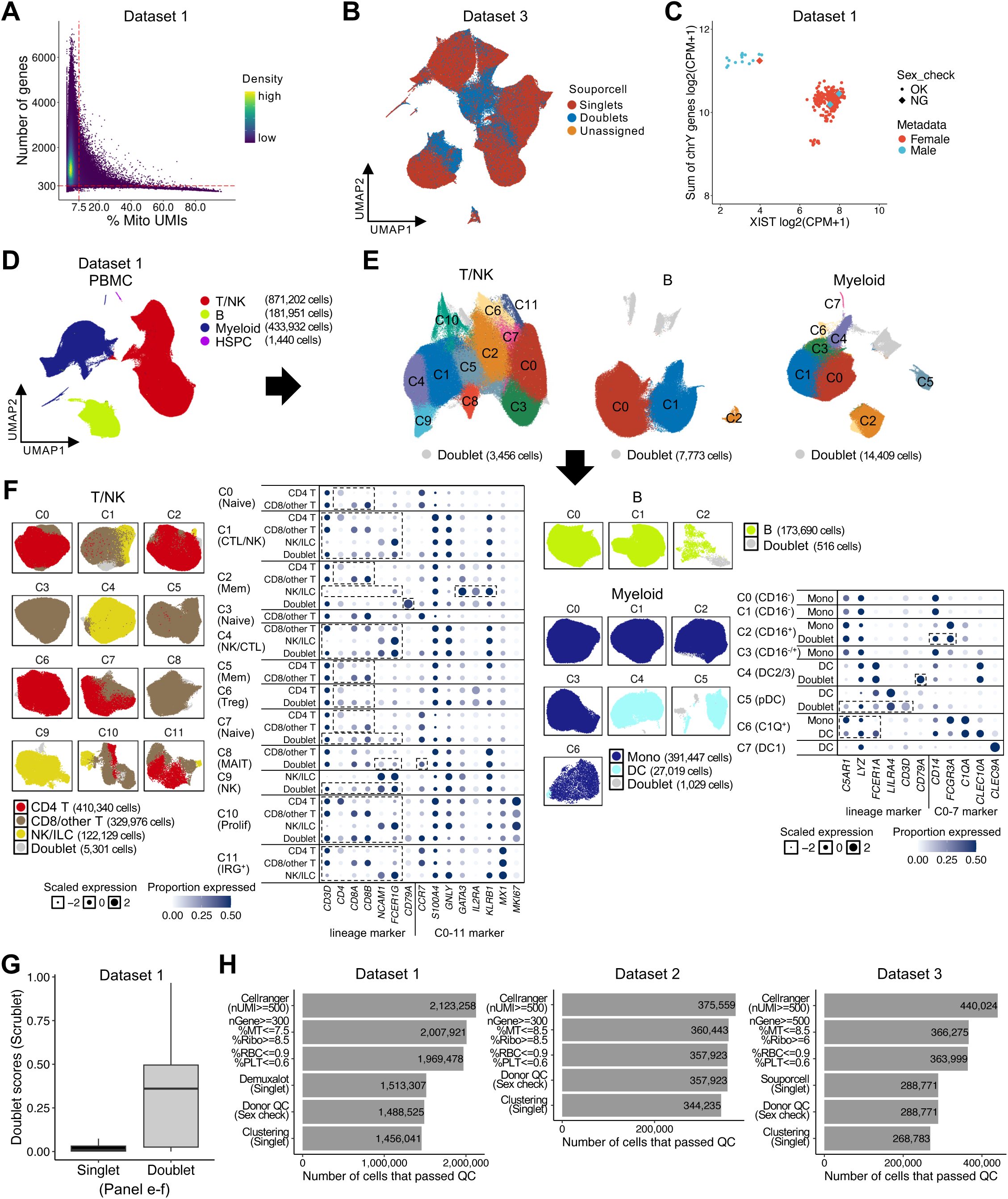
Quality control of datasets. **(A)** Density plot showing cell distributions based on the number of detected genes and the percentage of mitochondrial UMIs (%Mito) in dataset 1 (*N* = 269). **(B)** UMAP plot showing PBMCs in dataset 3 (*N* = 32), colored by Souporcell-based singlet/doublet/unassigned classification. **(C)** Scatter plot showing donor-level pseudobulk expression of XIST and Y-chromosome-related genes in dataset 1, colored by metadata-derived sex information. **(D)-(F)** Flowchart illustrating the strategy to partition PBMCs into seven lineages in dataset 1. In **(F)**, dot plots display scaled expression levels of lineage or subcluster marker genes, with color intensity indicating the proportion of expressing cells. **(G)** Boxplot comparing Scrublet-derived doublet scores between singlets and doublets identified in **(E)** and **(F)**. **(H)** Bar plots showing the number of cells retained after each QC step in datasets 1, 2, and 3.

**Fig. S2.**
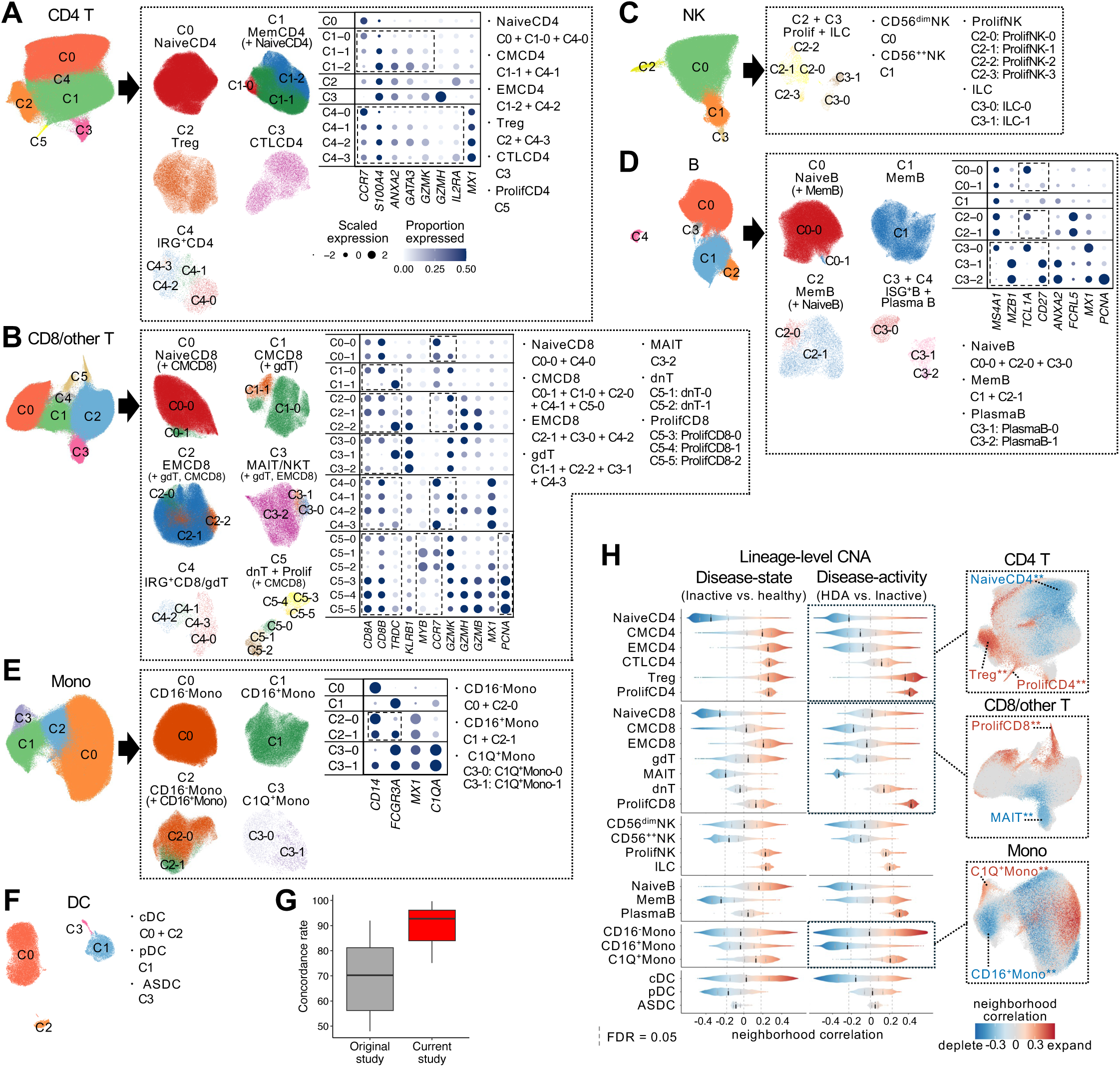
Definition of 123 robust cell states using fine-grained clustering. **(A-F)** Flowcharts illustrating the hierarchical separation of lineage cells into cell types in **(A)** CD4^+^ T cell (CD4 T), **(B)** CD8^+^ and unconventional T cell (CD8/other T), **(C)** natural killer cell (NK), **(D)** B cell, **(E)** monocyte (Mono), and **(F)** dendritic cell (DC) lineages (dataset 1; *N* = 269). The hematopoietic stem and progenitor cell (HSPC) lineage is omitted as it contains only a single cell type. Dot plots display scaled expression levels of cell-type marker genes, with color intensity representing the proportion of cells expressing each gene. **(G)** Box plot showing the concordance rates of cell-type-level annotation in the original and the current study with the predicted cell-type labels in the reference mapping onto the independent PBMC dataset. **(H) (left)** Distributions of neighborhood correlations for each cell type in disease-state (*n* = 137) or disease-activity (*n* = 61) comparisons (dataset 1, lineage-level-CNA). **(right)** UMAP plots in representative lineages. Cell types are highlighted if > 50% of neighborhoods are associated with phenotypes. See **table S2** for cell type abbreviations.

**Fig. S3.**
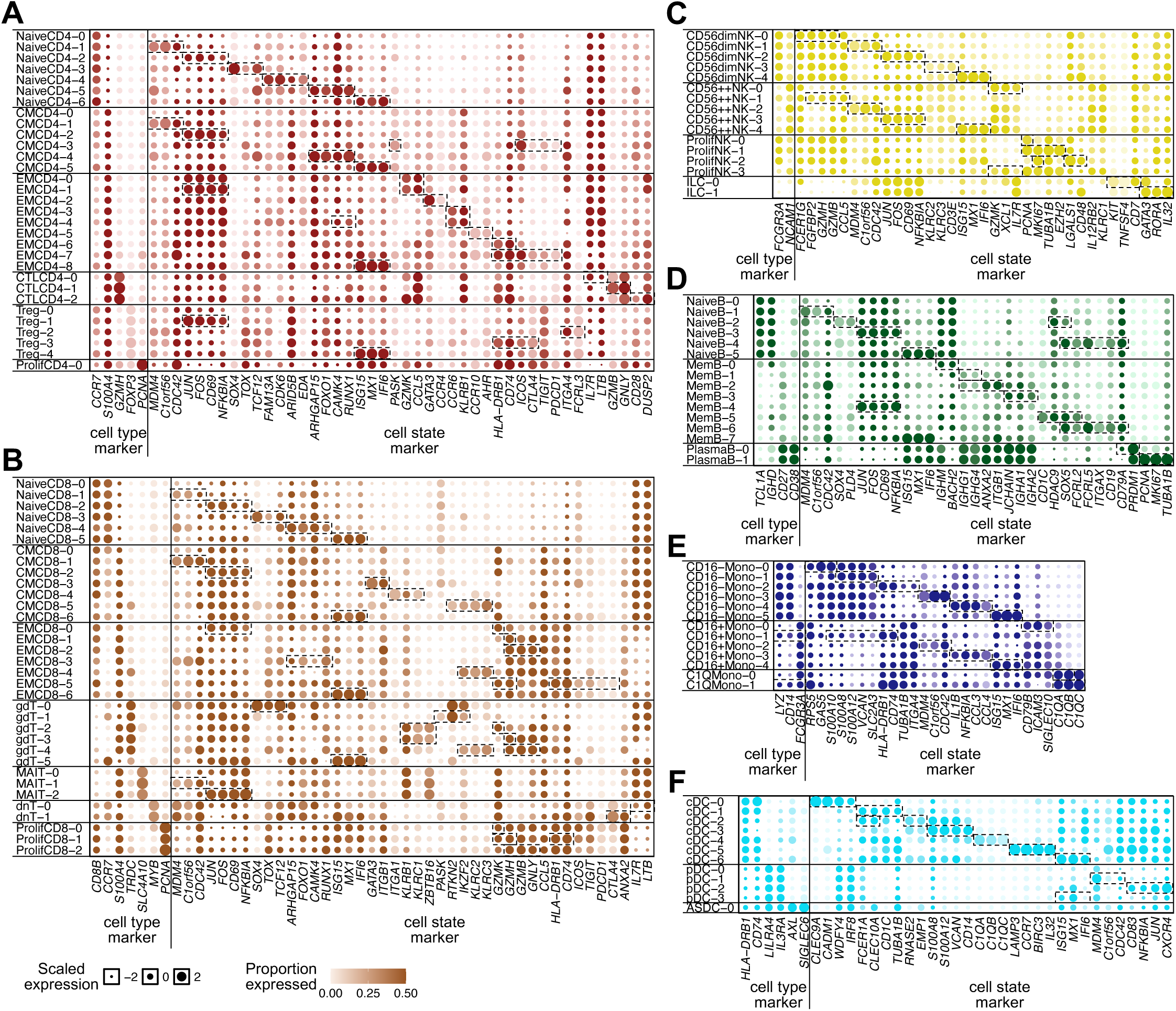
Marker gene expression across 123 cell states. **(A–F)** Dot plots showing scaled expression levels of cell-type and cell-state marker genes for each cell state in **(A)** CD4^+^ T cell (CD4 T), **(B)** CD8^+^ and unconventional T cell (CD8/other T), **(C)** natural killer cell (NK), **(D)** B cell, **(E)** monocyte (Mono), and **(F)** dendritic cell (DC) lineages (dataset 1; *N* = 269). Color intensity represents the proportion of cells expressing each gene.

**Fig. S4.**
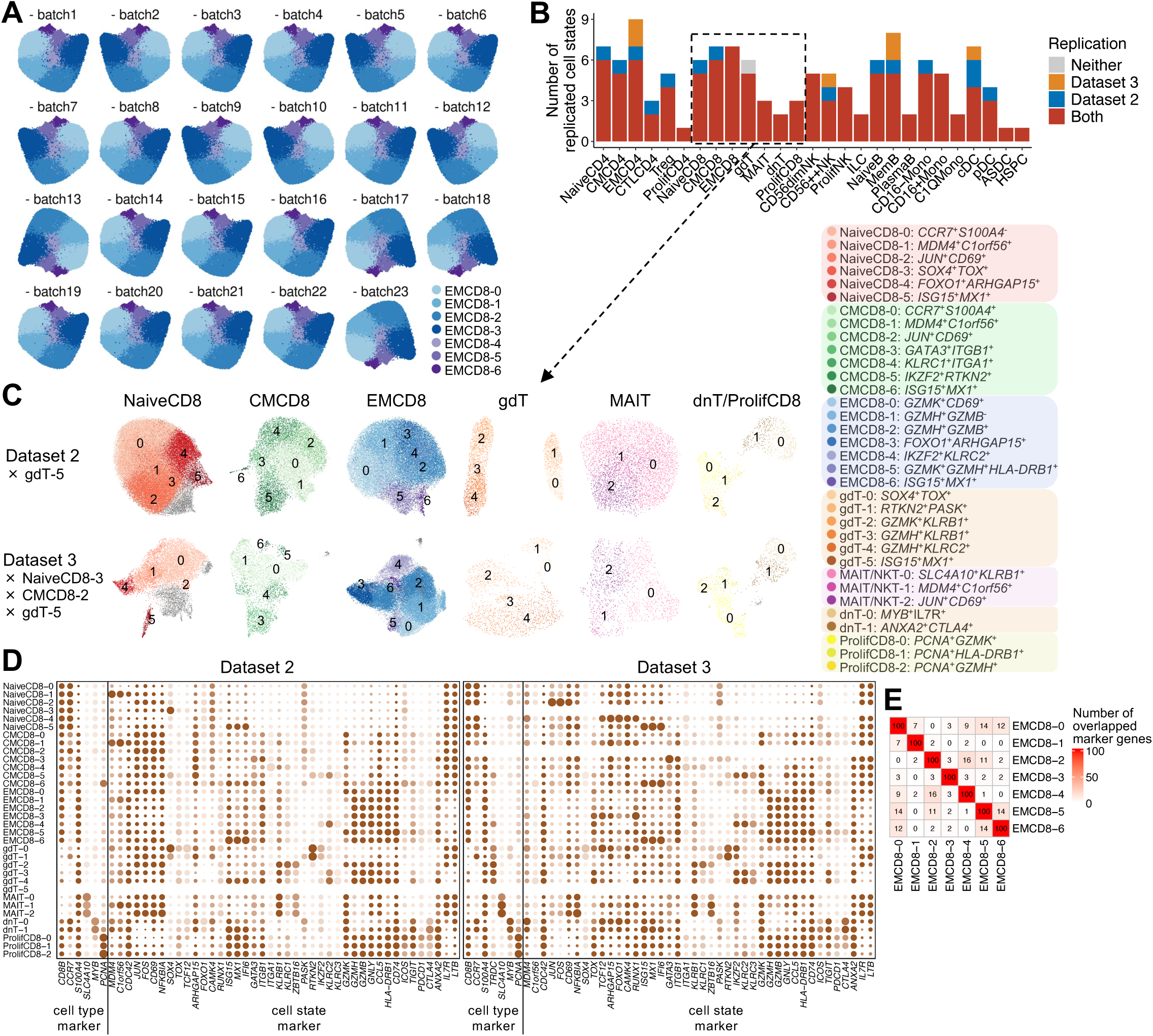
Robustness and replicability of 123 cell states. **(A)** UMAP plots of representative cell type (EMCD8), colored by cell states for each leave-one-batch-out condition. **(B)** Bar plots showing the number of cell states replicated in independent clustering of dataset 2 and/or 3. **(C)** UMAP plots of representative cell types in the CD8/other T lineage, colored by cell states in independent clustering of dataset 2 and 3. Cell states not replicated in each condition are listed to the left, and cell-state annotations are listed to the right. **(D)** Dot plots showing scaled expression levels of cell-type and cell-state marker genes for each cell state in the CD8/other T lineage from dataset 2 and 3. Color intensity represents the proportion of cells expressing each gene. **(E)** Heatmap showing the number of overlapping genes among the top 100 upregulated marker genes for each cell state in EMCD8.

**Fig. S5.**
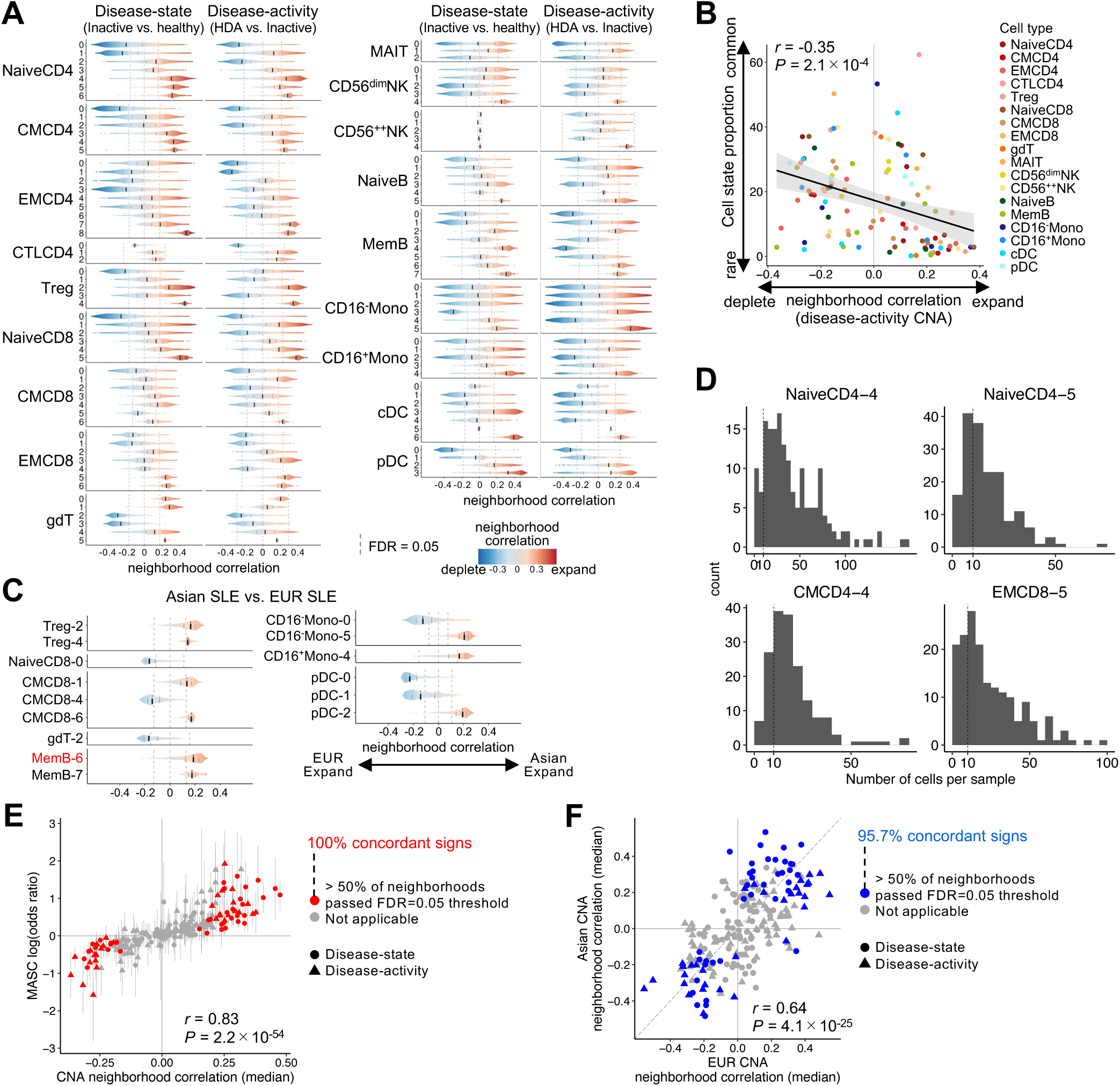
Mapping cell state compositional changes in SLE. **(A)** Distributions of neighborhood correlations for each cell state in disease-state (*n* = 137) or disease-activity (*n* = 61) comparisons (dataset 1, cell-type-level-CNA). **(B)** Scatter plot of median neighborhood correlation (disease-activity CNA) for each cell state vs. cell-state proportion within cell type. Shaded regions represent 95% confidence intervals. **(C)** Distributions of neighborhood correlations in Asian vs. EUR SLE comparisons. 15 cell states with > 50% of neighborhoods associated with phenotype (FDR < 0.05) are shown. **(D)** Histograms showing the number of cells per SLE sample for representative cell states expanded in SLE. **(E)** Scatter plot of median neighborhood correlation in CNA vs. effect size in MASC for each cell state in disease-state and activity comparisons. **(F)** Scatter plot of median neighborhood correlation of cell-type-level CNA in European vs. in Asian. In **(B)**, **(E)** and **(F)**, Pearson’s *r* and *P* values are indicated.

**Fig. S6.**
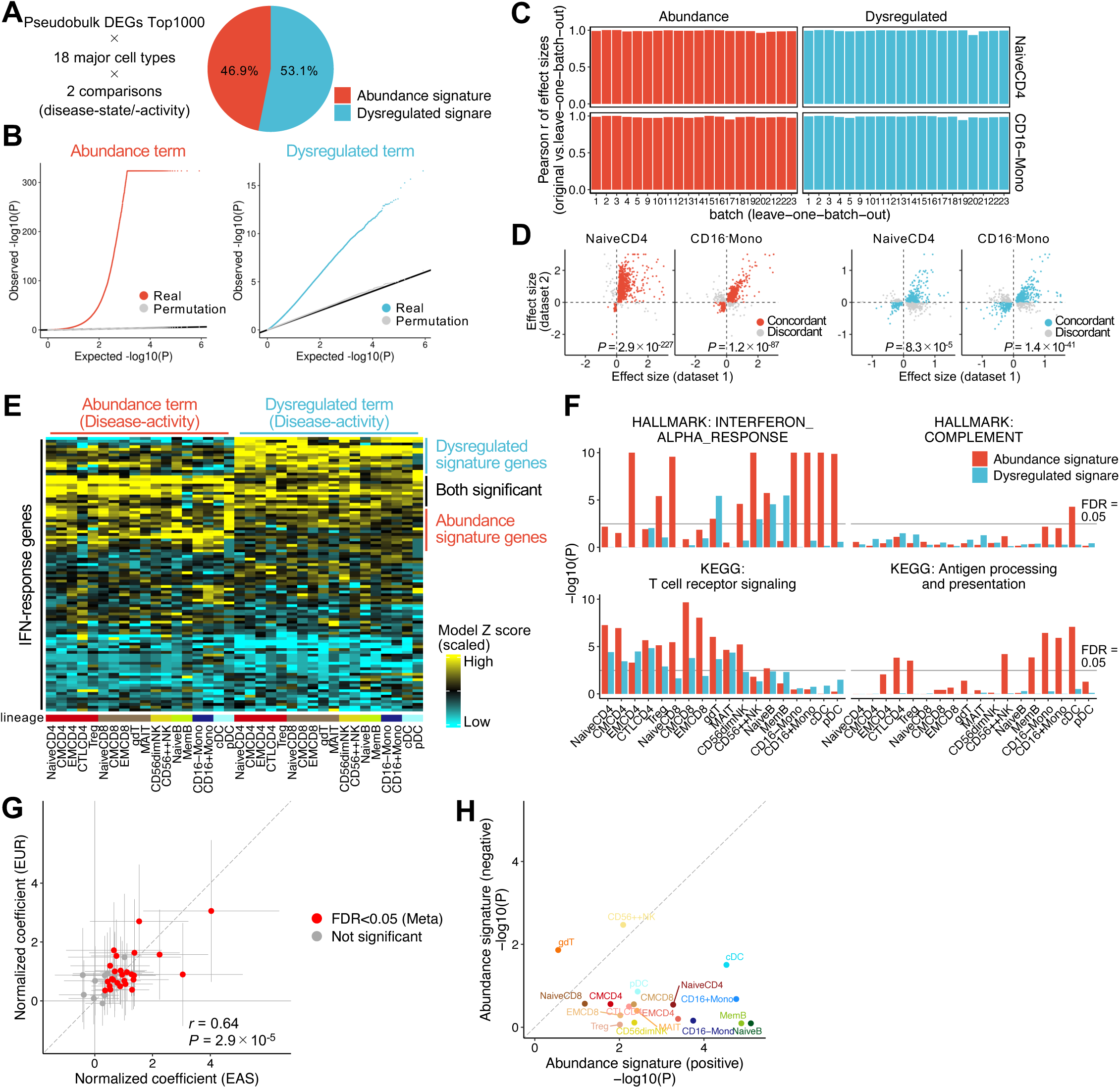
Statistical framework to dissect quantitative and qualitative changes within cell types. **(A)** Relative proportion of variance explained by abundance and dysregulated terms for the top 1,000 pseudobulk DEGs across all cell types and comparisons. **(B)** Quantile-quantile plots of observed vs. expected model *P* values for abundance and dysregulated terms across all cell types and comparisons. **(C)** Bar plot showing Pearson’s *r* of model effect sizes between the original vs. each leave-one-batch-out analysis for representative cell types. **(D)** Comparison of model effect sizes across genes between dataset 1 and 2. *P*, *P* values from one-sided binomial tests. **(E)** Heatmap of abundance and dysregulated term Z scores across 18 major cell types for 100 IFN-response genes (disease-activity comparison; *n* = 61). Gene order is based on hierarchical clustering of Z scores. **(F)** Bar plots showing the enrichment of representative pathways for each signature across cell types (disease-activity comparison). *P*, *P* values from one-sided Fisher’s exact tests. **(G)** Comparison of normalized coefficients in stratified LD score regression (S-LDSC) between EAS vs. EUR SLE-GWAS. Each dot represents a signature for a given cell type. Error bars represent 2×standard errors. Pearson’s *r* and *P* values are indicated. **(H)** Comparison of SLE risk variant enrichment around abundance signature genes with positive vs. negative model statistics for each cell type.

**Fig. S7.**
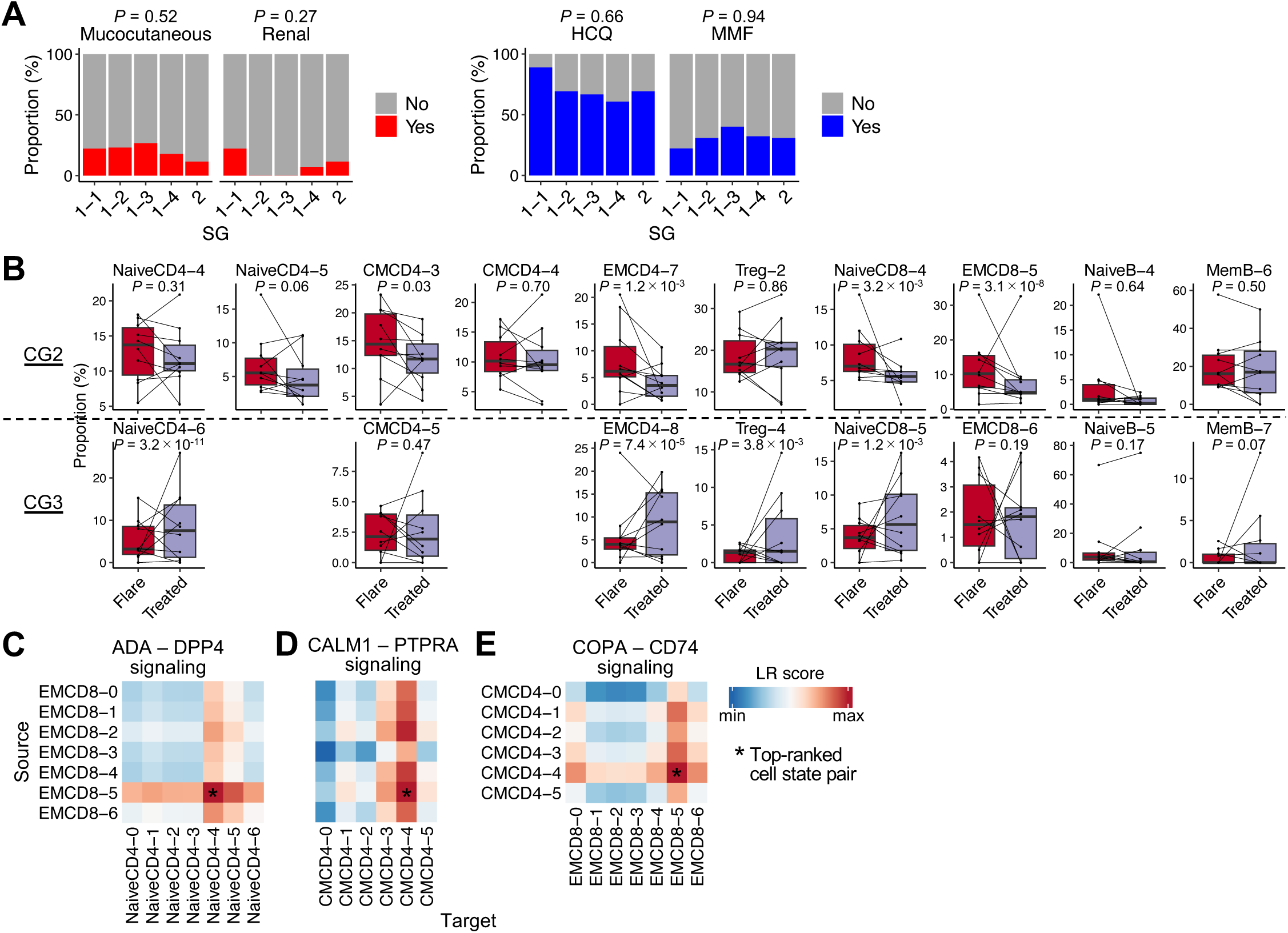
Characterization of aberrant immune networks between disease-relevant cell states. **(A)** Bar plot showing the frequencies of SLE samples with **(left)** mucocutaneous or renal activity and **(right)** hydroxychloroquine (HCQ) or mycofenolate mofetil (MMF) treatment for each sample group (SG). *P*, *P* values from two-sided Fisher’s exact tests. **(B)** Box plots comparing cell-state proportions between “Flare” and “Treated” status in **(top)** CG2 and **(bottom)** CG3 cell states. Samples from the same donor are connected by lines. *P*, *P* values from MASC (*n* = 10×2). **(C-E)** Heatmaps of LR scores for **(C)** ADA–DPP4 signaling from EMCD8 to NaiveCD4 cell states, **(D**) CALM1–PTPRA signaling from EMCD8 to CMCD4 cell states, and **(E)** COPA–CD74 signaling from CMCD4 to EMCD8 cell states.

**Fig. S8.**
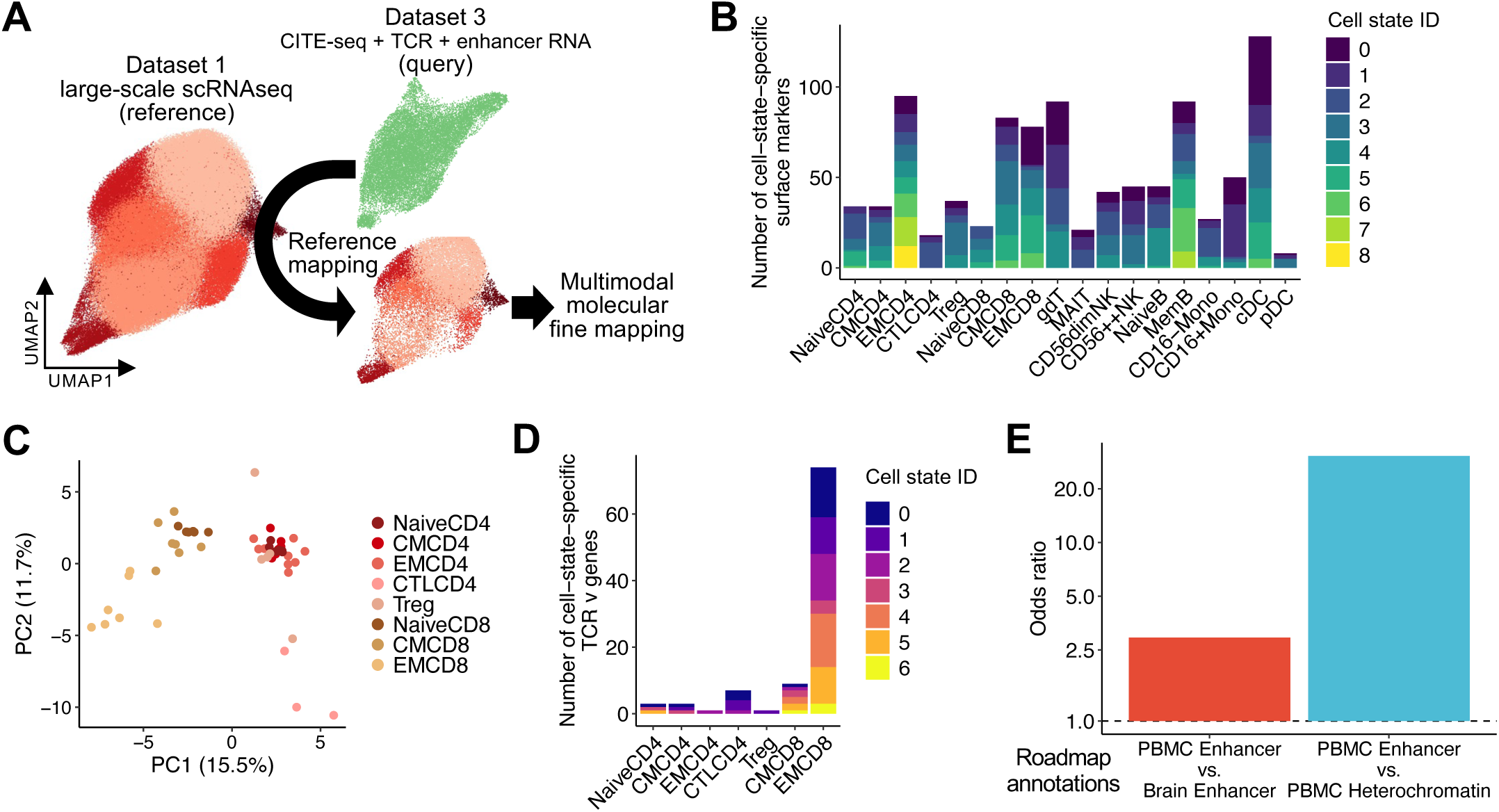
Multimodal molecular profiling across fine-grained cell states. **(A)** Schematic of the integration of multimodal dataset 3 (query; *N* = 32) into large scRNA-seq dataset 1 (reference; *N* = 269) through reference mapping. **(B)** Bar plot showing the number of cell-state-specific upregulated surface marker proteins. Colors represent cell-state IDs (e.g., NaiveCD4-0). **(C)** PCA plot of TCR V gene usage frequencies for each cell state, colored by cell type. **(D)** Bar plot showing the number of cell-state-specific (overrepresented) TCR V genes. **(E)** Bar plot showing the enrichment of called transcribed enhancers across all cell types for “PBMC enhancer” compared with **(left) “**Brain enhancer” and **(right) “**PBMC heterochromatin” in Roadmap Epigenomics project annotations.

**Fig. S9.**
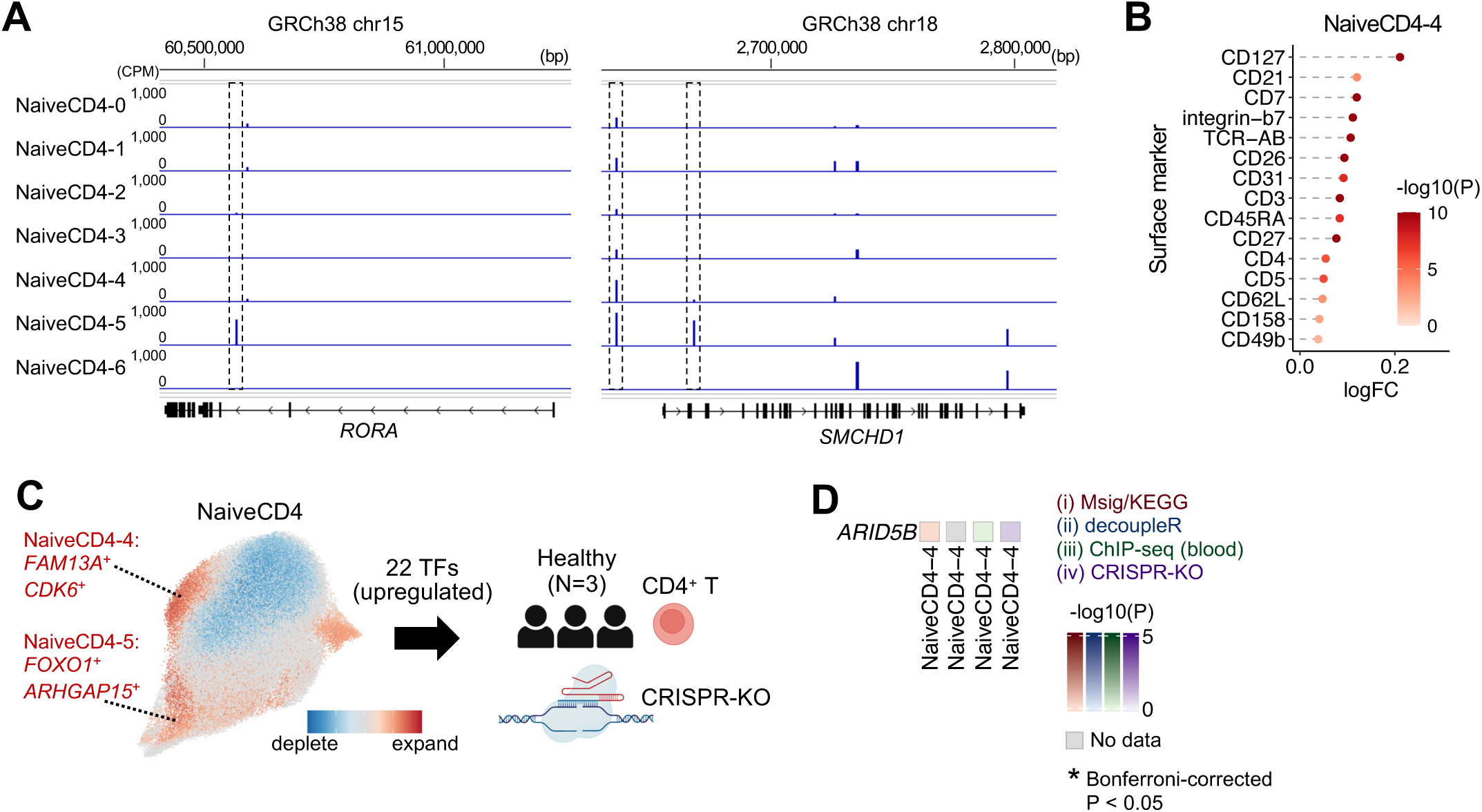
*FOXO1* is a candidate driver of the pathogenic CD4+ T cell state gene program. **(A)** Representative examples of cell-state-specific transcribed enhancers near upregulated marker genes in NaiveCD4-5. **(B)** Dot plot showing the logFC of top 15 upregulated surface markers for NaiveCD4-4. *P*, *P* values from Seurat FindMarkers. **(C)** Experiment design of the arrayed CRISPR-KO experiment in primary CD4^+^ T cells. **(D)** Heatmap showing - log_10_(enrichment *P*) values of *ARID5B* for NaiveCD4-4 signature genes based on (i) pathway/TF enrichment analyses, TF activity inference using (ii) decoupleR and (iii) blood-derived ChIP-seq data, and (iv) a CRISPR-KO experiment. *P*, *P* values from one-sided Fisher’s exact tests.

**Fig. S10.**
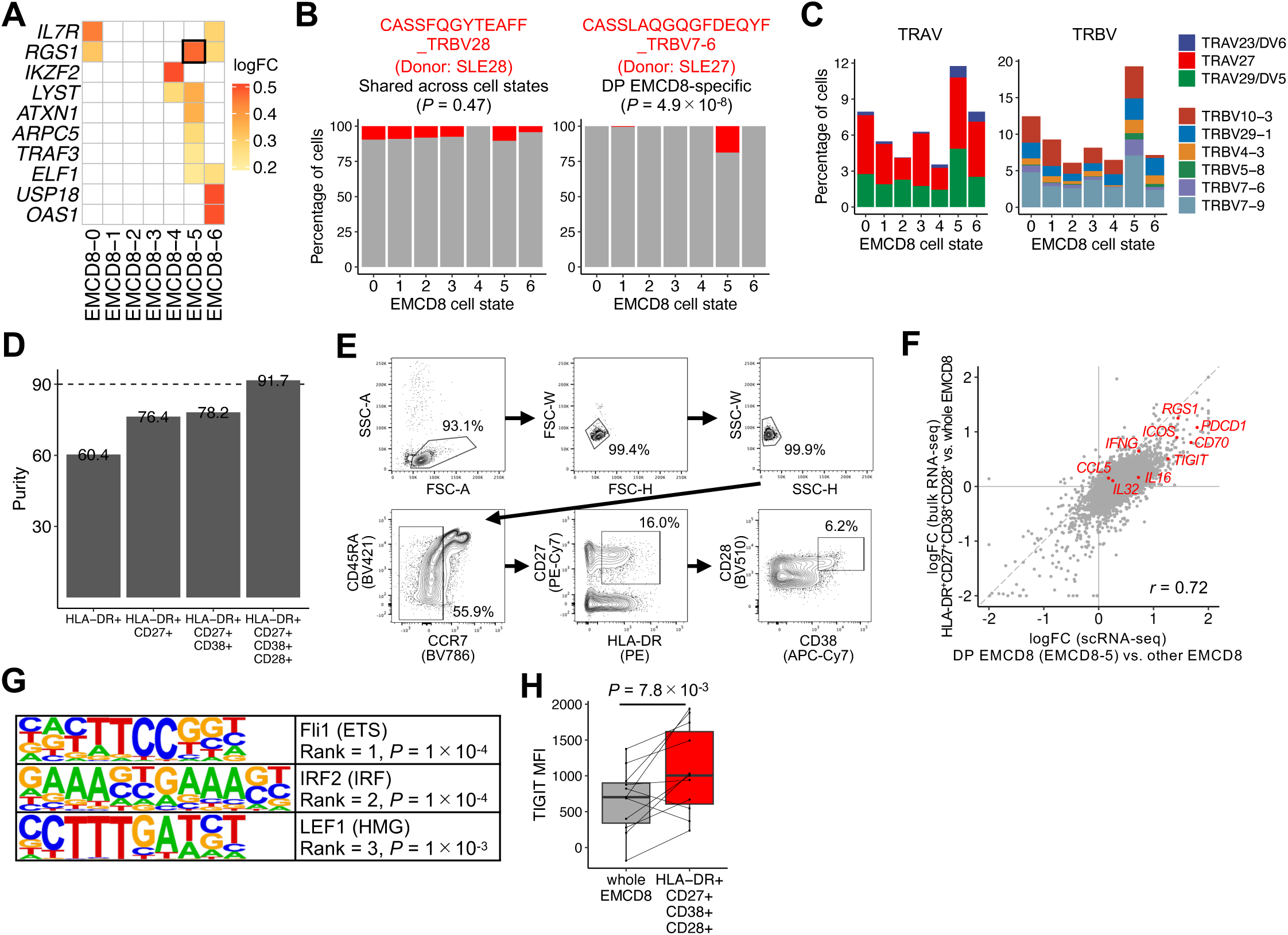
DP EMCD8 cells as a candidate pathogenic player in SLE. **(A)** Heatmap showing the logFC of GWAS-prioritized genes across EMCD8 cell states. Up to five genes with the highest logFC per cell state are shown. **(B)** Bar plots showing the percentage of **(left)** a representative clonotype shared across EMCD8 cell states and **(right)** a clonotype specific to EMCD8-5. *P*, *P* values from two-sided Fisher’s exact tests. **(C)** Bar plots showing the percentage of **(left)** three TRAV and **(right)** six TRBV gene usages positively skewed in EMCD8-5. **(D)** Bar plot showing the percentage of EMCD8-5 cells within each in-silico gate using dataset 3. **(E)** Gating strategy for defining HLA-DR^+^CD27^+^CD38^+^CD28^+^ EMCD8 cells in flowcytometry (FCM) and FACS. **(F)** Scatter plot comparing the scRNA-seq dataset 1 logFC (EMCD8-5 vs. other EMCD8 cells) and bulk RNA-seq logFC (HLA-DR^+^CD27^+^CD38^+^CD28^+^ vs. whole EMCD8 cells). **(G)** TF motif logos enriched for differentially accessible peaks in HLA-DR^+^CD27^+^CD38^+^CD28^+^ cells compared with whole EMCD8 cells. *P*, enrichment *P* values from HOMER. **(H) (left)** Boxplots comparing the mean fluorescence intensities (MFI) of TIGIT between HLA-DR^+^CD27^+^CD38^+^CD28^+^ and whole EMCD8 cells. Samples from the same donor are connected by lines. *P*, *P* values from linear mixed models.

## Methods

### Subjects

In this study, we utilized two publicly available 3’-end scRNA-seq datasets (datasets 1 and 2). For dataset 1, FASTQ files were downloaded from CELLxGENE (*5*) and raw genotype data were obtained from dbGaP (phs002812.v1.p1) with authorized access. Samples were recruited from the California Lupus Epidemiology Study (CLUES) cohort and the ImmVar Consortium. For dataset 2, count data were downloaded from dbGaP (phs002048.v1.p1) (*4*). For our newly generated 5’-end multimodal dataset (CITE-seq, TCR-seq, and enhancer RNA profiling; dataset 3) and additional FCM/FACS experiments, patients were recruited at Okayama University Hospital and Hospital of the University of the Occupational and Environmental Health under approvals by the Ethics Committees of RIKEN Center for Integrative Medical Sciences (RIKEN−Y-2025-006, 008). Written informed consent was obtained from all participants.

Across all datasets, SLE disease activity was categorized into four levels based on the SLE Disease Activity Index (SLEDAI) (*77*): i) inactive (SLEDAI = 0), ii) low (LDA; 1 ≤ SLEDAI ≤ 4), iii) moderate (MDA; 5 ≤ SLEDAI ≤ 9), and iv) high disease activity (HDA; SLEDAI ≥ 10) (**table S1**). In dataset 1, “flare” status was categorized into HDA.

### Sample processing and quality control

For dataset 3, single-cell suspensions of PBMCs from 32 SLE donors were processed using the 10x Genomics Chromium system. Gene expression (GEX), surface protein (ADT), and TCR (VDJ) libraries were prepared using the Chromium Next GEM Single Cell 5’ Kit v2 and TotalSeq™-C Human Universal Cocktail v1.0 (Biolegend, 399905). Eight libraries (16 samples pooled per library; four libraries per batch) were sequenced on an Illumina NovaSeq 6000 (150-bp paired-end).

FASTQ files for datasets 1 and 3 were processed using Cell Ranger v7.0.1. We excluded low-quality cells based on their distributions: (i) total unique molecular identifiers (UMIs) (< 500), (ii) number of genes (nGene), (iii) percentage of mitochondrial genes (% MT), or (iv) percentage of ribosomal protein genes (% Ribo) (dataset 1: nGene < 300, %MT > 7.5%, %Ribo < 8.5%; dataset 2: nGene < 300, %MT > 8.5%, %Ribo < 8.5%; dataset 3: nGene < 500, %MT > 8.5%, %Ribo < 6%; **fig. S1A**). Cells with high percentage of hemoglobin genes (> 0.9%; *HBB*, *HBA1*, *HBA2*, *HBD*) or platelet (> 0.6%; *PF4*, *PPBP*, *CAVIN2*, *GNG11*) signatures were also removed.

For dataset 1, samples were demultiplexed using Demuxalot (*78*) based on imputed genotypes (SHAPEIT2, Minimac3 r^2^ > 0.95) or Souporcell (*79*) for samples lacking reference genotypes. For dataset 3, Souporcell was used with 1000 Genomes Project common variants (minor allele frequency > 1%), followed by multiplex PCR and MiSeq sequencing of 20 target variants to link genotype clusters to specific individuals. Putative doublets from different genotypes were excluded (**fig. S1B**). In dataset 1,we excluded two samples with < 200 cells recovered and three samples showing sex discordance between metadata and pseudobulk expressions of *XIST* and Y chromosome genes (**fig. S1C**).

PBMCs were partitioned into seven lineages (CD4 T, CD8/other T, NK, B, Mono, DC, and HSPC) through iterative Louvain clustering (**fig. S1D–F**). Briefly, gene expression was normalized via SCTransform (Seurat v5.1.0) (*80*) and integrated using 3,000 highly variable genes (HVGs), followed by batch correction via Harmony (group.by.vars = “library”, “sample”) (*81*). Mitochondrial, Y chromosome, hemoglobin, platelet, and cell-cycle genes were excluded from HVGs. We used 30 Harmony-corrected PCs for UMAP visualization and shared nearest neighbor (SNN) graph construction. After initial separation into four major populations (T/NK, B, Myeloid, and HSPC), we performed recursive sub-clustering to maximize lineage purity based on marker gene expression (**fig. S1D–F**). Hyperparameters for each step are summarized in **table S3**. Sub-clusters representing remaining doublets with multiple lineage markers (**fig. S1F**) and higher Scrublet doublet scores (*82*) (**fig. S1G**) were excluded. After multi-step QC, we retained 1,456,041 (dataset 1), 344,235 (dataset 2), and 268,783 (dataset 3) high-quality PBMCs (**fig. S1H**).

### Definition of 123 fine-grained cell states

To identify granular cell states, we employed a three-layered iterative clustering strategy in dataset 1: lineage (layer 1), cell type (layer 2), and cell state (layer 3). Following initial lineage separation, we performed SCTransform normalization, HVG selection, PCA, Harmony batch correction, and Louvain clustering for each lineage using optimized parameters (**fig. S2A–F** and **table S3**). Sub-clustering was performed iteratively to ensure cell-type purity based on canonical marker expression. In a few instances (nine clusters), biologically redundant clusters were merged to define a final set of 123 cell states. To validate annotation accuracy, we performed reference mapping of dataset 1 onto an independent PBMC reference (Azimuth) (*19*) and compared concordance rates between our fine-grained annotations and the original labels (**fig. S2G**).

Cell-state-specific marker genes were identified using Seurat’s FindMarkers (v4.3.0) (min.pct = 0.05), comparing each state against all others within the same cell type (**fig. S3** and **table S4**). Upregulated markers (FDR < 0.05 and logFC > 0.1) were used for downstream pathway/TF enrichment analyses. To provide more intuitive differential expression values, we also reported logFC calculated with pseudocount.use = 0 in addition to the default Seurat values (**table S4**).

The robustness of the 123 cell states was assessed via leave-one-batch-out analysis, where each of the 23 experimental batches was systematically excluded, and the entire pipeline—from normalization to clustering—was re-executed. We titrated the resolution parameter (in increments of 0.05) to confirm that the original clusters were consistently replicated (**fig. S4A** and **table S5**). Furthermore, independent replication analyses were performed using dataset 2 and 3, where the entire pipeline was re-run from scratch for each dataset (**fig. S4B–C**). We confirmed that the resulting clusters exhibited marker gene profiles comparable to those in dataset 1 (**fig. S4D** and **table S5**).

### Co-varying neighborhood analysis

To identify single-cell-resolution transcriptional neighborhoods associated with clinical phenotypes, we applied co-varying neighborhood analysis (CNA v0.1.6) (*15*), to the 18 major cell types (> 5,000 cells each; **Fig. 3**) and six cell lineages (**fig. S2H**) in dataset 1. Following our established framework (*6*), we defined two comparison axes: (i) disease-state (97 healthy controls vs. 40 inactive SLE) and (ii) disease-activity (40 inactive vs. 21 high-activity [HDA] SLE).

Harmony-corrected PCs were used as inputs to infer the neighborhood abundance matrix (NAM), followed by PCA to obtain NAM-PCs. We tested the associations between phenotype labels and NAM-PCs using linear models, adjusting for age, sex, ancestry, and the number of cells per cell type. Given the substantial correlation between disease activity and age (*r* = -0.60) in dataset 1, age was excluded from covariates in disease-activity comparisons. For cell-type-level CNAs, significance was defined as a global permutation *P* < 0.05/36 (Bonferroni correction). CNA-derived neighborhood correlations (positive for expansion, negative for depletion) were aggregated for each cell state (**Fig. 3** and **table S8**). States were categorized as showing “remarkable expansion/depletion” if > 50% of constituent neighborhoods were associated with the phenotype at an empirical FDR < 0.05, and “moderate” if > 25%. We additionally performed CNA for (1) inactive vs. low-activity (LDA), (2) inactive vs. moderate-activity (MDA), and (3) Asian vs. European SLE (**Fig. 3E** and **fig. S5C**). MASC (*11*) was also applied to test cell-state abundance associations, adjusting for age, sex, and ancestry (**fig. S5E**).

For replication, CNA was applied to dataset 2 (**Fig. 3F**). Cells were projected into the dataset 1 latent space using Symphony (*53*), and Symphony-corrected PCs served as inputs. To ensure stable CNA performance (requiring ∼30 samples) (*15*), we compared 14 SLE (SLEDAI <= 2) vs. 17 healthy donors for disease-state, and 15 SLE (SLEDAI >= 6) vs. 14 SLE (SLEDAI <= 2) for disease-activity comparison.

### Statistical framework to dissect quantitative and qualitative changes

To decompose cell-type-level gene expression into quantitative and qualitative changes, we introduced a single-cell-level generalized linear mixed effect model (GLMM, **Fig. 4B** and **table S9**). For each gene, the model was fitted using a negative binomial distribution:

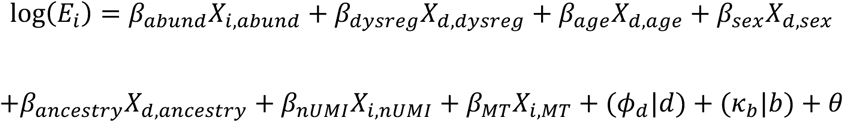

where *E* is the expression of the gene in cell *i*, *θ* is an intercept, and *β* coefficients represent fixed effects for cell *i*, donor *d*, or batch *b*. Donor and batch were modeled as random intercepts. The abundance signature term (*X_i,abund_*) represents neighborhood correlation values from cell-type-level CNA, while the dysregulated signature term (*X_d,dysreg_*) represents phenotype information (e.g., healthy vs. inactive SLE). The model was applied to genes passing low-expression filters (>=3 counts in >= 10% of samples) across 18 major cell types (> 5,000 cells) for disease-state and activity comparisons in dataset 1, using lme4 (*83*). Significance was determined via likelihood ratio tests (LRT) comparing models with and without the respective terms. ImmVar samples were excluded from disease-state comparisons. To quantify the contribution of each term to pseudobulk expression, we calculated Nagelkerke R^2^ values for the top 1,000 DEGs (identified via edgeR (*84*)) per cell type (**fig. S6A**).

Type I error calibration was assessed via permutation tests (randomly shuffling neighborhood correlations or phenotype labels; **fig. S6B**). Robustness was further validated through leave-one-batch-out analysis (**fig. S6C**) and independent replication in dataset 2, with effect-size replicability tested via one-sided binomial sign tests (**fig. S6D**). For 100 IFN response genes (IRGs) (*4*) (**table S11**), abundance and dysregulated Z-scores were used for hierarchical clustering (Ward’s method, Euclidean distance) to visualize patterns across cell types (**Fig. 4C–D** and **fig. S6E**).

### Pathway enrichment analysis and gene signature scores

To evaluate the enrichment of pathway annotations for abundance and dysregulated signature genes, we performed over-representation analysis (ORA) using the top 1,000 genes (by LRT P-values) for each signature term in each cell type. ORA was conducted using a one-sided Fisher’s exact test via clusterProfiler (*85*) (**Fig. 4E**, **fig. S6F**, and **table S12**), utilizing MSigDB Hallmark and KEGG pathway collections. To capture cell-type-specific biology, the union of the top 1,000 LRT P-value genes across all cell types was used as the background gene set.

For pathway analysis of cell-state-specific upregulated marker genes (FDR < 0.05, logFC > 0.1, Seurat FindMarkers), ORA was performed similarly (**Fig. 6A**, **Fig. 7A**, and **table S7**). The background for each cell type consisted of all genes passing the low-expression filter (>=3 counts in >= 10% of samples). To calculate classical exhaustion signature scores in EMCD8 and EMCD4 populations, an exhaustion-related gene set was curated from previous studies (*3*, *5*) (**table S20**). Gene expression was log-normalized and scaled, and the signature score for each cell was defined as the sum of the scaled expression values across the gene set (**Fig. 7B**).

### Stratified LD score regression analysis

To evaluate the enrichment of SLE heritability (all risk variants irrespective of effect size) around abundance and dysregulated signature genes, we performed stratified linkage disequilibrium score regression (S-LDSC) (*40*) using the sc-linker pipeline (*86*) (**Fig. 4F** and **fig. S6H**). We first calculated min-max-scaled gene scores based on the Z-scores of abundance and dysregulated terms for all tested genes in each cell type. LD scores were then estimated for East Asian (EAS) and European (EUR) populations using 1000 Genomes Project reference panels, with the scaled gene scores serving as annotation weights. In S-LDSC, we examined the enrichment of SLE heritability for common variants within 100-kb windows flanking the transcription start sites (TSS) of genes with positive Z-scores for both signatures, adjusting for the baseline model annotations v1.2 (*40*). Two large-scale SLE-GWAS summary statistics from EAS and EUR ancestries (*44*, *45*) were utilized. To account for differences in GWAS heritability, regression coefficients were normalized by the mean per-SNP heritability as previously described (*40*). Finally, a fixed-effect meta-analysis of the EAS and EUR results was performed using the inverse-variance weighting method based on the normalized coefficients and their standard errors (**fig. S6G**). We reported one-sided *P* values to test whether the regression coefficients were significantly positive.

### Localization of GWAS prioritized genes

To identify candidate causal genes for SLE-associated genetic variants (*P* < 5 × 10^−8^), we used the L2G (locus-to-gene) scores from the Open Targets Platform (*43*). We defined 124 genes with an L2G score > 0.5 as SLE-GWAS prioritized genes (**table S10**). For each cell state, we identified the overlap between these 124 prioritized genes and the cell-state-specific upregulated markers (FDR < 0.05, logFC > 0.2; Seurat FindMarkers) (**Fig. 6E** and **fig. S10A**).

### Sample group (SG) and cell-state group (CG) analysis

To investigate the clinical relevance of fine-grained cell states, we performed bi-hierarchical clustering (Ward’s method, Euclidean distance) of all samples in dataset 1 and cell states within 18 major cell types based on their proportions (**Fig. 5A**). Sample groups (SGs) and cell-state groups (CGs) were defined based on the resulting dendrograms. The frequencies of disease-activity categories, organ-specific activity, and treatment agents (**table S1**) were compared across SGs (**Fig. 5B** and **fig. S7A**). For longitudinal analysis in dataset 1, changes in cell-state proportions between paired “Flare” and “Treated” samples (*n* = 10 donors) were tested using MASC (*11*) (**Fig. 5C** and **fig. S7B**).

### Cell-cell interaction analysis

To infer the inflammatory networks across all cell states, we performed cell-cell interaction analysis using LIANA (46) with patemeters method = c(“logfc”, “sca”) and min_cells = 10. From the consensus set of 4,701 ligand–receptor (LR) interactions across all 123 × 123 cell-state pairs, we identified 1,451,770 potentially active signals (defined by sca.LRscore > 0.6, logfc.logfc_comb > -0.1, and expression proportions of >=1% for ligands and >=5% for receptors). To minimize redundancy, for each unique LR interaction within each of the 27 × 27 cell-type pairs, we prioritized the cell-state pair with the highest LR score as the representative cell-state-specific interaction (118,629 signals; **Fig. 4D–G**, **Fig. 6B**, **Fig. 7D**, and **fig. S7C–E**).

### Reference mapping and integration in LuPIN

To integrate the three single-cell datasets into the LuPIN framework, we performed reference mapping using Symphony (*53*), designating the large-scale dataset 1 as the reference and datasets 2 and 3 as queries (**fig. S8A**). For dataset 1, lineage- and cell-type-level reference objects were constructed based on the Harmony-integrated embeddings used in the fine-grained clustering. For the query datasets (2 and 3), PBMCs were first partitioned into seven lineages using the same pipeline as for dataset 1. Following normalization and scaling, cells from each lineage were projected into the corresponding lineage-level latent transcriptional space of dataset 1 (lineage-level reference mapping). Cell-type labels were then assigned to query cells using k-nearest neighbor classifiers (k = 5) based on the dataset 1 reference. This procedure was repeated at the cell-type level (cell-type-level reference mapping) to annotate each cell in datasets 2 and 3 with the best-predicted cell-state labels.

### Surface protein analysis

Following reference mapping of the multimodal dataset 3, we investigated cell-state-specific surface protein profiles. For dataset 3, 130 surface proteins and seven isotype controls were profiled using the TotalSeq™-C Human Universal Cocktail v1.0 (Biolegend, 399905). Protein expression for each cell was normalized using a centered-log ratio (CLR) transformation across all 130 proteins. Cell-state-specific surface markers were identified using Seurat FindMarkers, comparing each cell state against all others within the same cell type. Proteins with FDR <0.05 and logFC >0.2 were defined as cell-state-specific upregulated surface markers (**fig. S8B** and **table S14**).

### TCR repertoire analysis

In dataset 3, we utilized TCR UMI counts for cells meeting the following criteria: (i) passed GEX-based QC; (ii) annotated as NaiveCD4, CMCD4, EMCD4, CTLCD4, Treg, NaiveCD8, CMCD8, or EMCD8 via reference mapping; and (iii) passed Cell Ranger TCR quality filters. To identify cell-state-specific V gene usage, frequencies were compared for each state against all others within the same cell type using two-sided Fisher’s exact tests (**Fig. 6C**, **fig. S8D**, **S10C**, and **table S15**). Among all 45 TRAV and 49 TRBV genes, we tested V genes passing a low expression filter (>= 10 total counts per cell type).

TCR clonal expansion in EMCD8-5 (DP EMCD8) was assessed by quantifying clonotypes across SLE donors (**Fig. 7C**). Clonotypes were defined based on the combination of the CDR3 amino acid sequence and the V gene of the β-chain. To identify DP EMCD8-specific clonotypes, frequencies were compared between states within each donor using two-sided Fisher’s exact tests (**fig. S10B** and **table S21**). Known cognate epitopes for each clonotype were identified using the VDJdb browser (**table S21**).

### Transcribed enhancer analysis

To detect transcribed enhancers in dataset 3, we employed the ReapTEC pipeline (*18*). 10x Genomics libraries were sequenced on an Illumina NovaSeq 6000 with 150-bp paired-end reads for both Read 1 and Read 2, ensuring the capture of transcription start sites (TSS) within the 5’ regions. Reads were mapped using STARsolo to identify those containing unencoded G. After deduplication, reads were assigned to specific cell types based on barcodes. TSS peaks were generated by merging TSSs within 10 bp, and bidirectional transcribed candidate enhancers were identified using the developers’ scripts, with promoter regions masked. To validate these enhancers, we tested their enrichment across Roadmap Epigenomics core 15-state reference annotations (*57*) for PBMC-derived enhancers (6_EnhG and 7_Enh in E062), compared with (i) PBMC heterochromatin (9_Het in E062) and (ii) brain hippocampus middle enhancers (6_EnhG and 7_Enh in E071) using two-sided Fisher’s exact tests (**fig. S8E**).

Cell-state-specific transcribed enhancers were identified using an entropy-based specificity score (*18*). We first filtered for enhancers with >=5 reads per cell type, aggregated reads across donors, and calculated entropy and specificity scores for each enhancer within each cell type. Enhancers with a specificity score > 0.7 were defined as cell-state-specific (**table S16**).

### Trajectory analysis

To infer the developmental trajectory across the CD4^+^ T cell lineage, we employed Monocle3 (*64*) with default parameters (**Fig. 6G**). For pseudotime estimation, the root was defined as a cell within the SOX4^+^TOX^+^ NaiveCD4-3 state. This selection was based on recent evidence that these cells represent recent thymic emigrants (RTEs), the most immature population within the peripheral naive T cell compartment (*22*).

### Upstream TF activity inference

To identify transcriptional regulators driving disease-relevant CD4^+^ T cell states, we integrated three complementary approaches (**Fig. 6F** and **fig. S9D**): (i) over-representation analysis (ORA) of TF targets, (ii) TF activity inference via decoupleR (*62*) and (iii) target enrichment analysis using ChIP-Atlas (*63*). Candidates TFs were selected from those upregulated (logFC > 0.3) in NaiveCD4-5 (n = 22) and NaiveCD4-4 (n = 1, *ARID5B*). ORA was conducted using a one-sided Fisher’s exact test via clusterProfiler (*85*) for MSigDB C3 and KEGG collections (**table S17**), using signature genes (FDR < 0.05, logFC > 0.1; Seurat FindMarkers) from NaiveCD4-4, NaiveCD4-5, and CMCD4-4. For decoupleR, TF activities were inferred using the ulm method based on DoRothEA regulons and FindMarkers logFC values. For the ChIP-Atlas-based analysis, we utilized ChIP-seq datasets restricted to blood tissues (**table S17**). Promoter regions were defined as -5kb to +2kb from the TSS. TF-target gene annotations were constructed by intersecting ChIP-seq peaks with these promoter regions. Enrichment of NaiveCD4-4/5, and CMCD4-4 signature genes within these TF-target sets was evaluated using one-sided Fisher’s exact tests.

### Arrayed CRISPR-KO experiment

To validate the identified transcriptional regulators, we performed an arrayed CRISPR knock-out (KO) experiment targeting 22 candidate TFs (**Upstream TF activity inference**; **fig. S9C**). For each TF, two independent single guide RNAs (sgRNAs) (*87*) were synthesized (IDT; **table S18**) and complexed with recombinant Cas9 (IDT, 1081058) to form ribonucleoprotein (RNP) complexes. Two control samples (four sgRNAs) were prepared for each donor. Editing and frameshift efficiency for each sgRNA was confirmed via Sanger sequencing (on median 79.7% and 68.7%, respectively; **table S18**).

Primary CD4^+^ T cells from three healthy donors were isolated from fresh PBMCs using CD4^+^ T Cell Isolation Kit (Miltenyi, 130-096-533), expanded for 2 days using Dynabeads Human T-Activator CD3/CD28 (Thermo, DB11132) in complete CTS OpTmizer T-Cell Expansion SFM (Thermo Fisher Scientific, A1048501) supplemented with IL-2 (Peprotech, 200-02), IL-7 (Peprotech, 200-07), and IL-15 (Peprotech, 200-15). Seven days after activation, cells were electroporated using the 4D-Nucleofector (Lonza, program EH-140) with 1 × 10⁶ cells per reaction and subsequently cultured in triplicate. Five days post-transfection, total RNA was isolated using MagMAX-96 Total RNA Isolation Kit (Thermo Fisher Scientific, AM1830) and libraries were prepared using the RamDA-seq method (*88*). Sequencing was performed on an Illumina NovaSeq 6000 (150bp paired-end).

For RNA-seq analysis, adaptors were trimmed using Cutadapt and reads with < 50 bp were removed. Reads were mapped to GRCh38 using STAR (*89*) and quantified via HTSeq in unstranded mode. After excluding three low-quality samples (unique reads < 4 × 10^6^), TMM-normalized counts (*84*) were fitted to a negative binomial GLMM using lme4 (*83*):

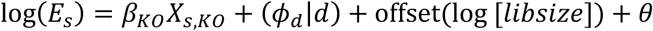

where *E* is gene expression in sample *s*, *θ* is an intercept, *β_KO_* is the fixed effect for KO status, and Donor (*d*) is a random intercept. Significance was determined via LRT. Enrichment of downregulated genes (FDR < 0.05; **table S19**) within NaiveCD4-4/5 and CMCD-4 signature genes was tested using one-sided Fisher’s exact tests (**Fig. 6F**).

### FCM and FACS experiments on EMCD8 cells

To characterize EMCD8-5 (DP EMCD8) cells, we performed functional assays using FACSMelody (BD Biosciences) based on the four surface markers with the highest logFC values (HLA-DR, CD27, CD38, and CD28; **Fig. 7E–F** and **table S14**). Two panels were prepared: Panel 1 for surface markers (PD-1, TIGIT) and Panel 2 for intracellular IFN-γ (**table S22**).

For FACS, CD8^+^ T cells were isolated from six SLE donors using the CD8^+^ T Cell Isolation Kit (Miltenyi, 130-096-495), blocked with anti-FcγR antibodies (Thermo Fisher Scientific), and stained with Panel 1. Target EMCD8 (CCR7^−^HLA-DR^+^CD27^+^CD38^+^CD28^+^) and total CCR7^−^ EMCD8 cells (control) were sorted (5,000 cells per sample; **fig. S10E**) for bulk RNA-seq (SMART-seq, Takara Bio 634772), ATAC-seq, and UNIChro-seq. ATAC-seq tagmentation used TDE1 (Illumina 20034198) and 1% digitonin (Promega G944A). Purified tagmented DNA was amplified for 15–18 cycles using KAPA HiFi ReadyMix (Nippon Genetics KK2602) and Nextera XT Index Kit v2 and size-selected (180–1000 bp) using AMPure XP beads. UNIChro-seq libraries were prepared with custom primers to quantify chromatin accessibility (*68*) at five target *IFNG* and one reference *UBE2D2* loci (**table S24**) and sequenced on Miseq (illumina) using a custom R1 primer (5’- ACACTCTTTCCCTACACGACGCTCTTCAGGTCT).

RNA-seq data were analyzed using the same pipeline as the **Arrayed CRISPR-KO experiment**, with the following GLMM:

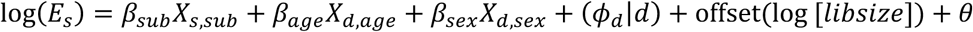

where *β_sub_* is the fixed effect for cell subsets. ATAC-seq and UNIChro-seq reads were mapped using Bowtie2 (*90*), with low-quality (q < 30), duplicated (Picard), and ENCODE-blacklisted reads removed. MACS2 (*91*) was used for peak calling. Differentially accessible regions (DARs) were identified using the same GLMM framework as RNA-seq. Motif enrichment for DARs (FDR < 0.1, logFC >0) was performed with HOMER (**fig. S10G**). For UNIChro-seq, we extracted UMI counts at target loci as previously described (*68*). Accessibility at five IFNG loci relative to a UBE2D2 reference was evaluated using a logistic mixed-effect model, regressing cell status (target/control EMCD8) by genomic region (target/reference) with donor as a random effect. Genomic tracks were visualized using Integrative Genomics Viewer (**Fig. 7G**).

For FCM, CD8^+^ T cells from 11 SLE donors were stained with Panel 1 (surface) or Panel 2 (intracellular). For Panel 2, cells were stimulated with PMA (50 ng/ml, MP Biomedicals, 16561-29-8) and ionomycin (1 μg/ml) for four hours, with GolgiStop (BD Biosciences, 554724) added after the first hour. Intracellular staining was performed using the Cytofix/Cytoperm kit (BD Biosciences, 554714). Mean fluorescence intensities (MFI) were compared between target and total EMCD8 cells using linear mixed models (fixed effects: age, sex; random effect: donor; **Fig. 7H** and **fig. S10H**). All data were analyzed using FlowJo (BD Biosciences).

